# Habitat loss in the restricted range of the endemic Ghanaian cichlid *Limbochromis robertsi*

**DOI:** 10.1101/2019.12.16.877282

**Authors:** A. Lamboj, O. Lucanus, P. Osei Darko, J.P Arroyo-Mora, M Kalacska

**Affiliations:** Department of Integrative Zoology, University of Vienna, Austria; Applied Remote Sensing Laboratory, Department of Geography, McGill University, Montreal QC, Canada; Flight Research Laboratory, National Research Council of Canada, Ottawa, ON. Canada

**Keywords:** agricultural expansion, deforestation, freshwater fish, gold mining, land use land cover change (LUCC), remote sensing, satellite imagery

## Abstract

Remote sensing, through satellite image analysis has become an integral and invaluable tool to inform biodiversity conservation and monitoring of habitat degradation and restoration over time. Despite the disproportionately high levels of biodiversity loss in freshwater ecosystems worldwide, ichthyofauna are commonly overlooked in favor of other keystone species. Freshwater fish, as indicators of overall aquatic ecosystem health can also be indicators of larger scale problems within an ecosystem. If endemic and specialized fishes are at risk, the forest and landscape around their habitat is also undergoing change. As a case study demonstrating the utility of multi-temporal, multi-resolution satellite imagery, we examined deforestation and forest fragmentation around the Atewa Forest Reserve, south eastern Ghana. Within small creeks, *Limbochromis robertsi*, a unique freshwater cichlid with an extremely limited distribution range can be found. Historically, the land cover in the area has undergone substantial deforestation for agriculture and artisanal small-scale mining, primarily for gold. We found deforestation accelerated along with increased forest fragmentation in the 2014 – 2017 period with the majority of the forest loss along the river and creek banks due to small-scale mining operations and increased agriculture. Field visits indicate a decrease in the total population by approximately 90% from the early 1990s to 2018. We illustrate the benefits of determining landscape metrics from local scale remote sensing studies as proxies to assess the decline of endemic species with restricted ranges, whose habitat characteristics and the subsequent pressures they face require detailed analysis at fine temporal and spatial scales not captured by global or continental scale datasets.

## REMOTE SENSING IN THE FORM OF SATELLITE IMAGE EARTH OBSERVATION PLAYS AN INCREASINGLY IMPORTANT ROLE IN SPECIES CONSERVATION (ROSE ET AL. 2015)

Since the early pioneering studies illustrating its utility for quantifying and mapping deforestation (e.g. Skole and Tucker 1993, Westman et al. 1989) and monitoring biodiversity (e.g. Case 1992, Stom and Estes 1993), remote sensing has become an integral aspect of biodiversity conservation worldwide (e.g. O’Connor et al. 2015, Paganini et al. 2016, Proenca et al. 2017, Vihervaara et al. 2017) detailing in most cases the rapid decline in habitat extent and quality (Pimm et al. 2001). As such, satellite image analysis provides an unbiased historical assessment of land cover change, current estimates of land cover extent– and serves as a monitoring tool. Following the rapid increase in the number of Earth Observation satellites from both national and international efforts (Tatem et al. 2008) over the last 20 years (e.g. Rapideye, Sentinel 2, Planet Dove nano-satellite constellation, etc.) and a variety of products (e.g. forest cover, albedo, land surface temperature, etc), assessments of biodiversity decline for endemic species can benefit from the use of these fine grained multi-temporal data. It is key for conservation practitioners to understand the correct use (and limitations) of these products and/or platforms, because they form the basis for extracted/modeled ecosystem variables and in many cases are used for decision making and policy development (de Leeuw et al. 2010, Congalton et al. 2014, Perez-Hoyos et al. 2017, Garcia-Alvarez et al. 2019, Martinez-Fernandez et al. 2019).

Despite freshwater ecosystems being highly threatened by anthropogenic activities and climate change, and subject to disproportionately high biodiversity loss worldwide (Ravenga et al. 2005, Geist 2011, Sundermann et al. 2013), freshwater ichthyofauna is often overlooked in biodiversity conservation. Increased advocacy for conservation in the terrestrial and marine environments (e.g. Hannah et al. 2002) has not yet had the same impact in freshwater ecosystems. While global scale databases and studies are beneficial for understanding anthropogenic drivers of land use and land cover change (LUCC) affecting populations at country or continental scales, endemic species with restricted ranges require studies at local scales which are not necessarily captured by global datasets (spatially or temporally). Such local (or basin) level studies can inform the implementation of conservation policies and measure progress in decreasing the rapid decline in habitat extent and quality for freshwater species (Ravenga et al. 2005) as a complement to the need for increased cyberinfrastructure for mapping biodiversity (e.g. Wheeler at al. 2012).

Artisanal small-scale mining (ASM and gold mining: ASGM) is practiced in many parts of the developing world including Ghana. The activity predates several centuries and continues today serving as an important activity sustaining the livelihood of many (Bansah et al. 2016; Hilson 2001). Apart from the contribution of ASGM to the national gold production (Hilson 2001), it is estimated that over ten percent of the Ghanaian population depends on ASGM for their livelihood (Bansah et al. 2018; Fritz et al. 2017); Ghana is one of top five gold producing countries in Africa from ASGM (Seccatore et al. 2014). The sector comprises licensed and unlicensed (i.e. “galamsey”) small-scale miners who use labor intensive extraction methods ranging from rudimentary to advanced machinery (Hilson 2001). Some miners seek placer gold deposits from river beds while others excavate ore bearing sediments from the subsurface producing large piles of sand which are further washed to extract the gold. Washing the sand with water and mercury is a common gold extraction process utilized by ASGM which poses a major threat to water bodies in many mining communities (Hilson 2001). It was estimated that in 2010, 37% of mercury emissions worldwide were from ASGM (UNEP 2013) with 1-2 g mercury lost per g of gold produced (Green et al. 2019). Over the years, the ASGM sector in Ghana has been heavily criticized for water and land pollution (Afum and Owusu 2016, Bansah et al. 2018), disturbance of flora and fauna and major flooding events in the rainy season (Nyame and Grant 2012). Ghanaian society has increasingly become interested in the discourse on socio-economic benefits and environmental risks prompting the government to ban all ASGM related activities in early 2017 and to initiate steps to formalize the sector (Meijer et al. 2018). Some reclamation work has been conducted following the ban and there are sites undergoing natural revegetation. Extensive economic development and population increase has led to LUCC with loss of natural vegetation cover to agriculture, illegal logging, urbanization and mining activities (Coulter et al. 2016; Hackman et al. 2017; Ntiamoa-Baidu et al. 2000). Ghana is estimated to have lost 80% of its forest cover throughout the twentieth century due to logging followed by slash and burn agriculture. It is also one of the first countries to report extinction of a primate, the red colobus monkey (*Procolobus badius waldroni*) due to destruction of its habitat (Ministry of Environment Science Technology and Innovation 2016, Oates et al. 2000). A land cover assessment for 2015 indicated that over 50 percent of the country is agriculture dominated with dense forests occupying only eight percent (mainly in forest reserves) (Hackman et al. 2017). In addition to natural barriers such as waterfalls, freshwater river and creek fragmentation is increasingly driven by anthropogenic factors, including pollution, LUCC and the introduction of non-native species (or expansion of the ranges of competitive native species) (Best, 2019).

LUCC resulting in loss of Riparian forest or water contamination from mining or agricultural runoff (Brito et al. 2018; Guo et al. 2015; Nobrega et al. 2018) have deleterious effects on clear water forest species such as *L. robertsi*. Environmental sex determination (ESD) has been shown in several fish including a West African cichlid, where a decrease in pH resulted in male biasing conditions (Reddon and Hurd 2013). Elevated water temperature is also an ESD factor resulting in a sex-ratio skewed towards the male phenotype (Abozaid et al. 2012; Baroiller et al. 2009; Bezault et al. 2007; Conover and Kynard 1981; Romer and Beisenherz 1996; Santi et al. 2016). Deforestation driven stream warming has further been shown to lead to reduction in fish body size (Ilha et al. 2018). In addition, exposure to increased photoperiod and the light spectrum from artificial lights at night, surrounding human development, disrupts freshwater and Riparian ecosystems and fish larval development (Boeuf and Le Bail 1999; Fugere et al. 2018; Perkin et al. 2011; Villamizar et al. 2009). Increase in benthic algal biomass has been shown to be positively correlated to lack of shade in streams and increased nutrient content (Fugere et al. 2018; Mebane et al. 2014; Zongo et al. 2017). Decline in freshwater river and stream quality fragments habitats (Fuller et al, 2015) and reduces the fish population density including a risk of local extinction (Brito et al. 2018).

In this study we used a combination of well-established satellite image analysis techniques for imagery with different temporal and spatial scales to assess changes in the landscape (i.e. LUCC) due to increased deforestation and mining activities. Baseline fragmentation statistics were used as a proxy to explain the observed decline of the endemic cichlid *Limbochromis robertsi* (van den Audenaerde and Loiselle 1971) in the Birim river basin around the Atewa Forest Reserve (Eastern region) in Ghana (Fig. 1). Multi-temporal semi-structured field observations over a 27 year period were carried out to document the presence/absence and abundance of the species. Furthermore, we discuss socio-economic aspects to account for the forest decline in the region. The creeks *L. robertsi* is known to inhabit were historically surrounded by forest but have undergone substantial degradation of the forested areas (Fig. 2). The multi-resolution and multi-temporal integration of high-resolution satellite imagery (2-3 m) with moderate resolution Landsat (30 m) data provides a detailed blueprint of the habitat degradation and urgent need for conservation. Effects of forest fragmentation, including the loss of area, increase in isolation, and greater exposure to LUCC have been shown to have long-term negative impacts on the structure and function of ecosystems (Haddad et al. 2015). We contribute to the growing body of knowledge on how these factors, well established in terrestrial environments, also impact freshwater aquatic habitats (Fuller et al. 2015). While our focus is on one species as a case study example, the methodology is widely applicable to other endemic and restricted range species worldwide.

**FIGURE 1.**
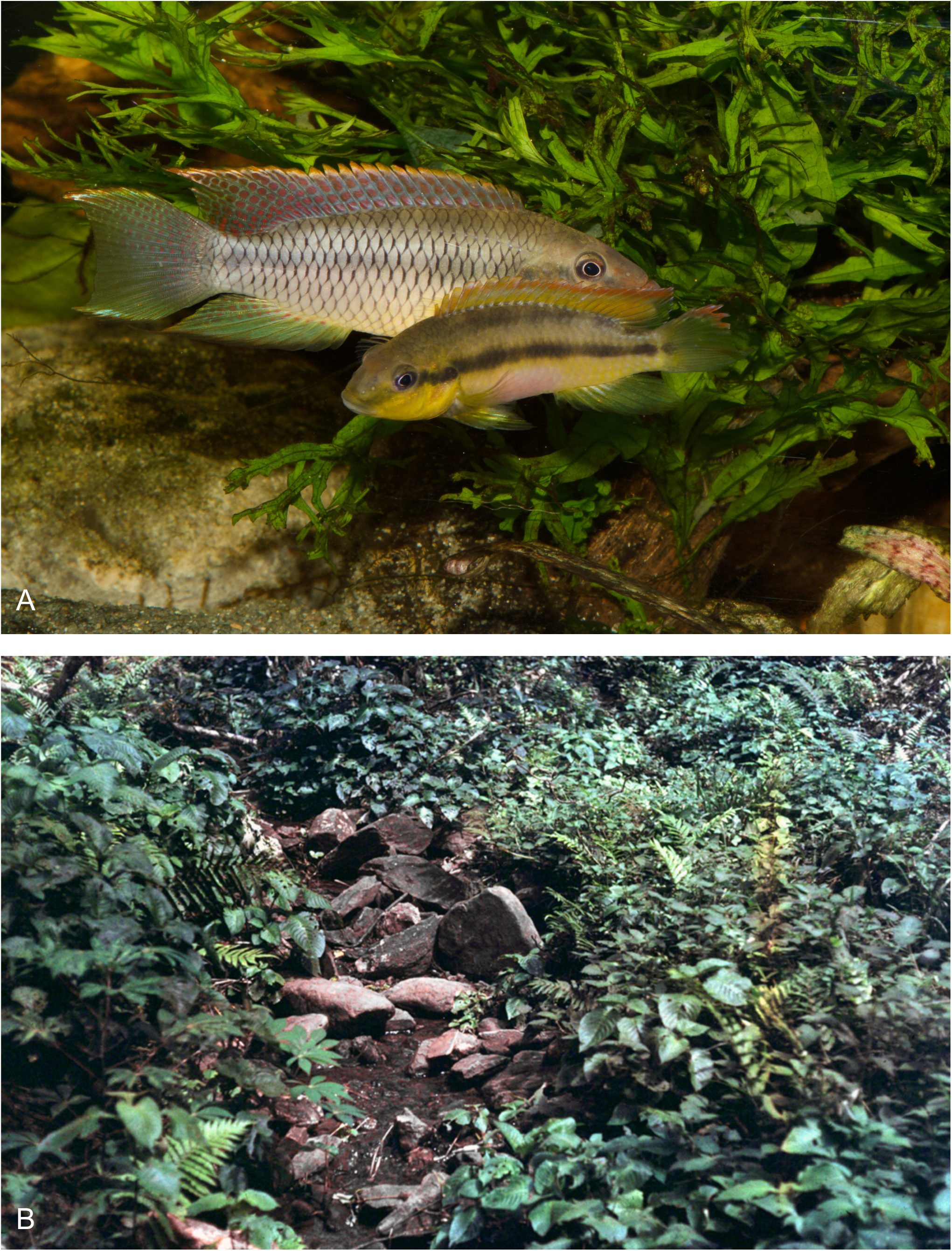
A) *Limbochromis robertsi* pair. The male is the larger individual. This photograph shows the species in breeding colour (photograph by A. Lamboj); B) Type locality of *L. robertsi*, region of Asiakwa, north of Kibi, Ghana. Photograph from 1966 (P. Loiselle).

**FIGURE 2.**
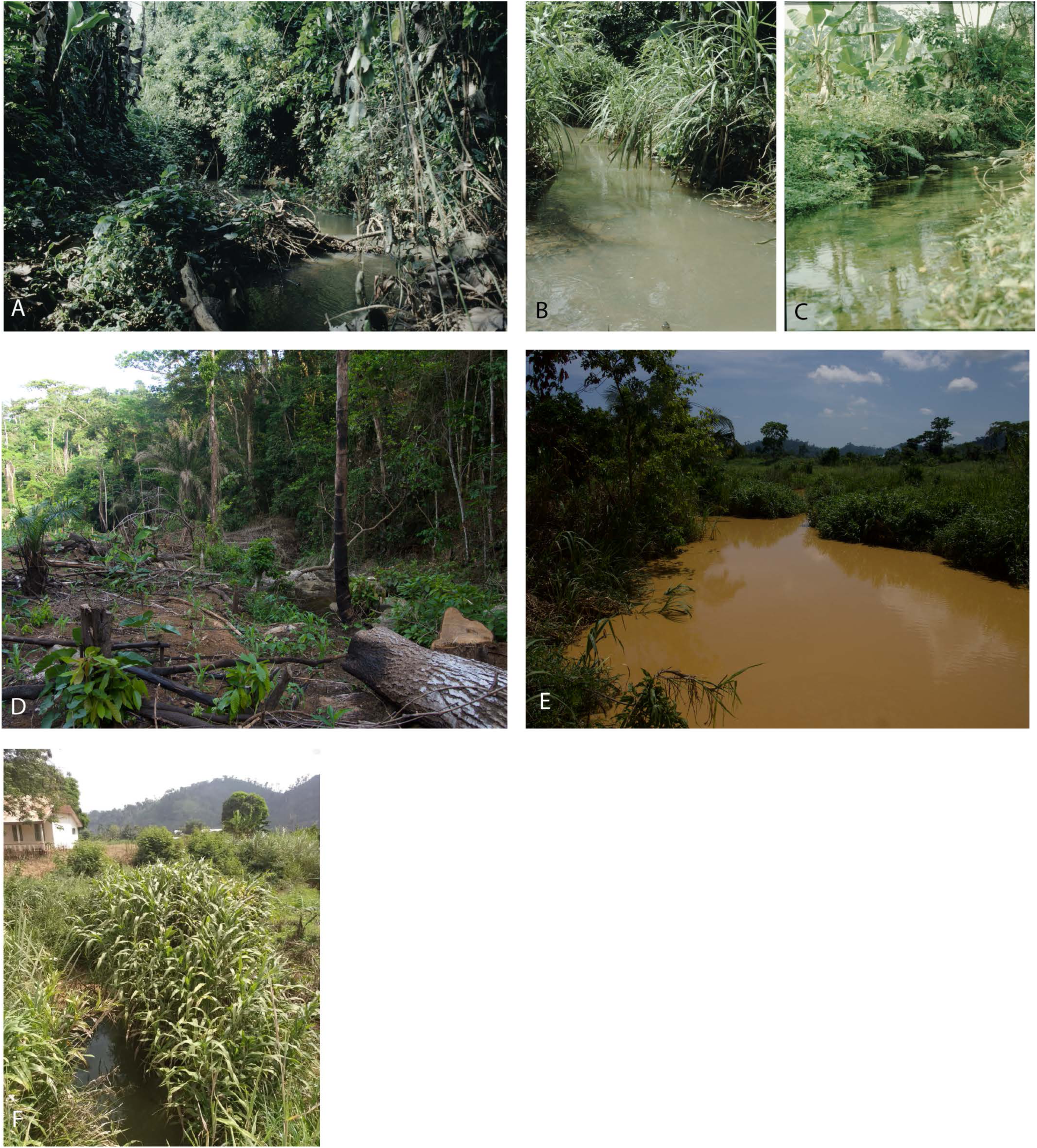
A) Black Krensen creek, 1991; B,C) Birim river, 1991; D) Black Krensen creek 2016; E) Birim river 2016; F) Black Krensen creek 2018. Photographs by A. Lamboj and E. Gyekety.

## METHODS

### CASE STUDY SPECIES

*Limbochromis robertsi*, the only member of its genus, and one of few vertebrates endemic to Ghana, is a unique freshwater fish (*Chromidotilapiine* cichlid) (Fig. 1A). It is primarily a cave breeding species with both parents involved in the brood care of the young, but unlike any other species of African cichlid it is able to switch its breeding strategy. Females may choose to pick up the young and raise them as a larvophile mouthbrooder, even if the pair stays together in their territory defended by the male (Lamboj 2006). While mouth brooding cichlids are not uncommon, *L. robertsi* is the only species known to use either breeding strategy in raising the offspring. It also has a special phylogenetic position within the group of *Chromidotilapiine* cichlids. This group has, in general, a wide distribution in West and Central Africa, ranging from Guinee in the east to the Congo River basin in the west (Lamboj 2004). Recent work based on molecular data suggests *Chromdotilapia schoutedeni* is the closest relative, a species from the upper Congo River (Kisangani region) (Schwarzer et al. 2014), making this species even more unique and of high scientific interest.

*L. robertsi* has an extremely limited distribution range; it is only known from the area around Kibi (also referred to as Kyebi) in Eastern Ghana. There, it is restricted to the middle elevations of small hill creeks (Fig. 1B, 2A, D, F) which drain into the Birim river (Fig. 2 B,C, E). In the early 1990s a second, very small population was observed from the western part of the country, in the Ankasa region of Ghana (U. Schliewenn pers. comm.). *L. robertsi* and *Chromidotilapia g. guntheri*, the only *Chromidotilapiine* cichlids occurring in Ghana, are not sympatric in the same rivers or creeks. In contrast to *L. robertsi*, *C. g. guntheri* prefers larger rivers in the lowlands and is widespread along the Niger river from Guinea to Nigeria. The two species are also readily differentiated: *L. robersi* is more elongate, males have a bifurcated caudal fin, yellow cheeks, reddish dots on the cheeks and on the ventral side of the body as well on the dorsal, caudal and anal fins. Females have horizontal red-silver-red bands on the dorsal fin, yellowish cheeks, silvery and red bands on the upper half of the caudal fin. In contrast, *C. g. guntheri* is deeper bodied, males are uniformly brown with red dots on the fins. Females have a silver dorsal fin, often with black dots and a reddish belly. *C. g. guntheri* is also a larger fish.

### STUDY AREA

Forest cover change was assessed in a 1389.5 km^2^ area of the Birim river basin around the Atewa Forest Reserve, Eastern region (Akwapim Togo Mountains ecoregion) (Fig. 3). The protected Atewa Forest Reserve was designated a Global Significant Biodiversity Area in 1939 and is one of only two reserves with upland evergreen forest in Ghana (Meijer et al. 2018). The Birim river is one of the main tributaries of the Pra river, a primary freshwater resource in the country. Over its length of approximately 181 km, the Birim crosses several large communities with a total population estimated at ∼2.6 million inhabitants (Ghana Statistical Survey 2012). The entire Birim river basin has been affected by ASM operations (Afum and Owusu 2016).

**FIGURE 3.**
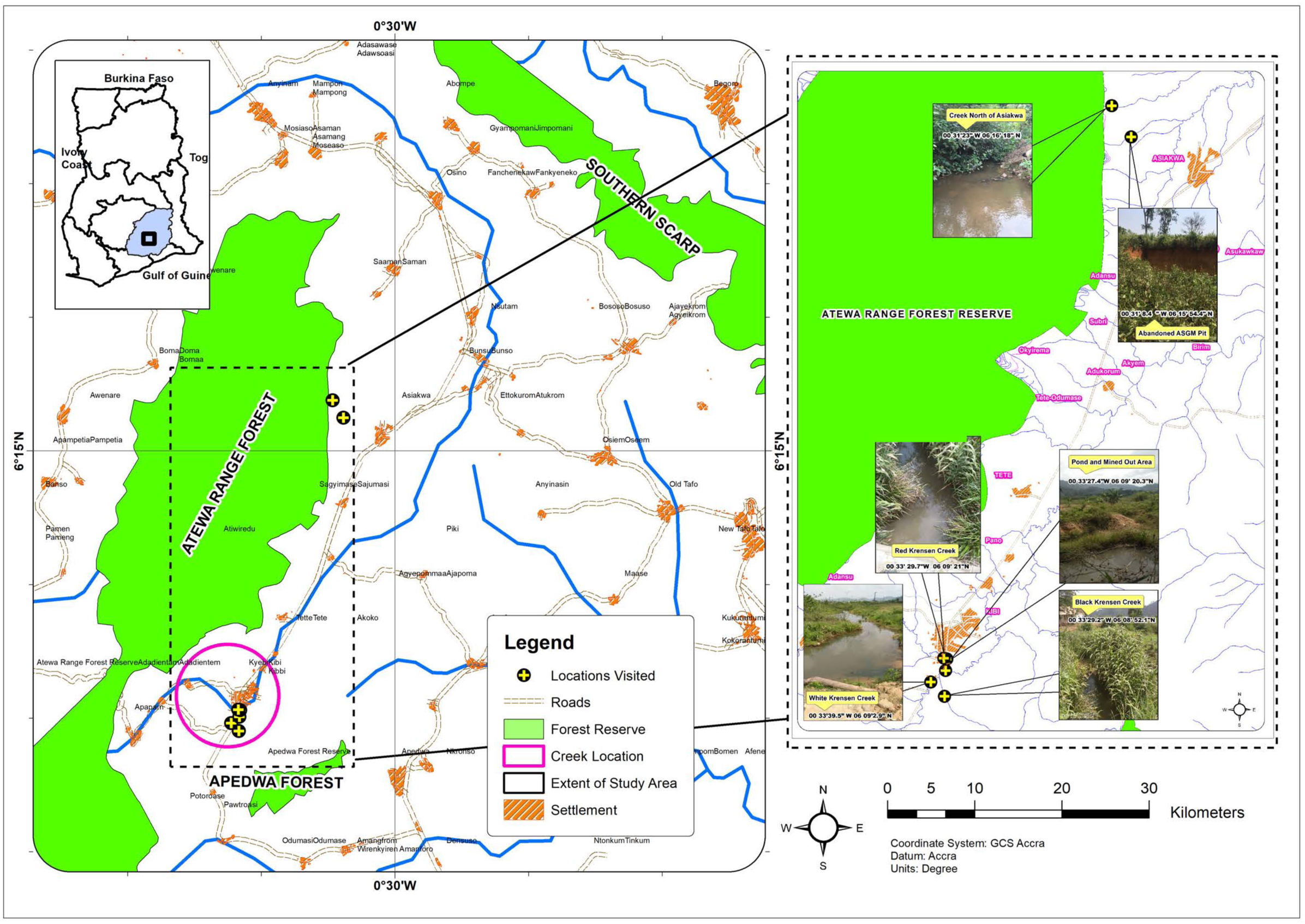
Map of the 1389.5 km^2^ study area of the Birim river basin around the Atewa forest reserve. Inset shows the location of the Black, Red and White Krensen creeks south of Kibi where populations of *L. robertsi* were observed in the early 1990s as well as the type locality and creeks visited in 2019.

### REMOTE SENSING DATA

To assess deforestation in the area (1389.5 km^2^) around the Atewa Forest Reserve, we used eight “Collection 1, Tier 1” radiometrically calibrated Landsat satellite images downloaded from the USGS Earth Explorer (Table S1). These images were acquired from 1986 to 2017. Landsat images have a 30 m pixel size for the multispectral bands. Landsat Thematic Mapper 5 (TM5) has five bands spanning the blue to shortwave infrared wavelengths. Landsat 7 Enhanced Thematic Mapper Plus (ETM+) has an additional shortwave infrared band (6 multispectral bands total), and Landsat 8 Operational Land Imagery (OLI) also has a coastal/aerosol band (7 multispectral bands total). Images were atmospherically corrected within CLASlite v3.2 to surface reflectance (Asner et al. 2009). This processing step is fundamentally important to remove the scattering and absorption effects from the atmosphere as much as possible to ensure the images are comparable over time and for an accurate classification. For a 25 km^2^ area near the collection site of *L. robertsi* south of Kibi we purchased high resolution satellite imagery (1968 – 2018) (Table S2, Fig. 4). These images range from a single panchromatic band for the declassified 70 mm film reconnaissance satellite photograph (1968) to 4 multispectral bands (blue, green, red, near infrared) for the contemporary 2-3 m pixel size IKONOS and Pleides 1A/1B satellite images.

**FIGURE 4.**
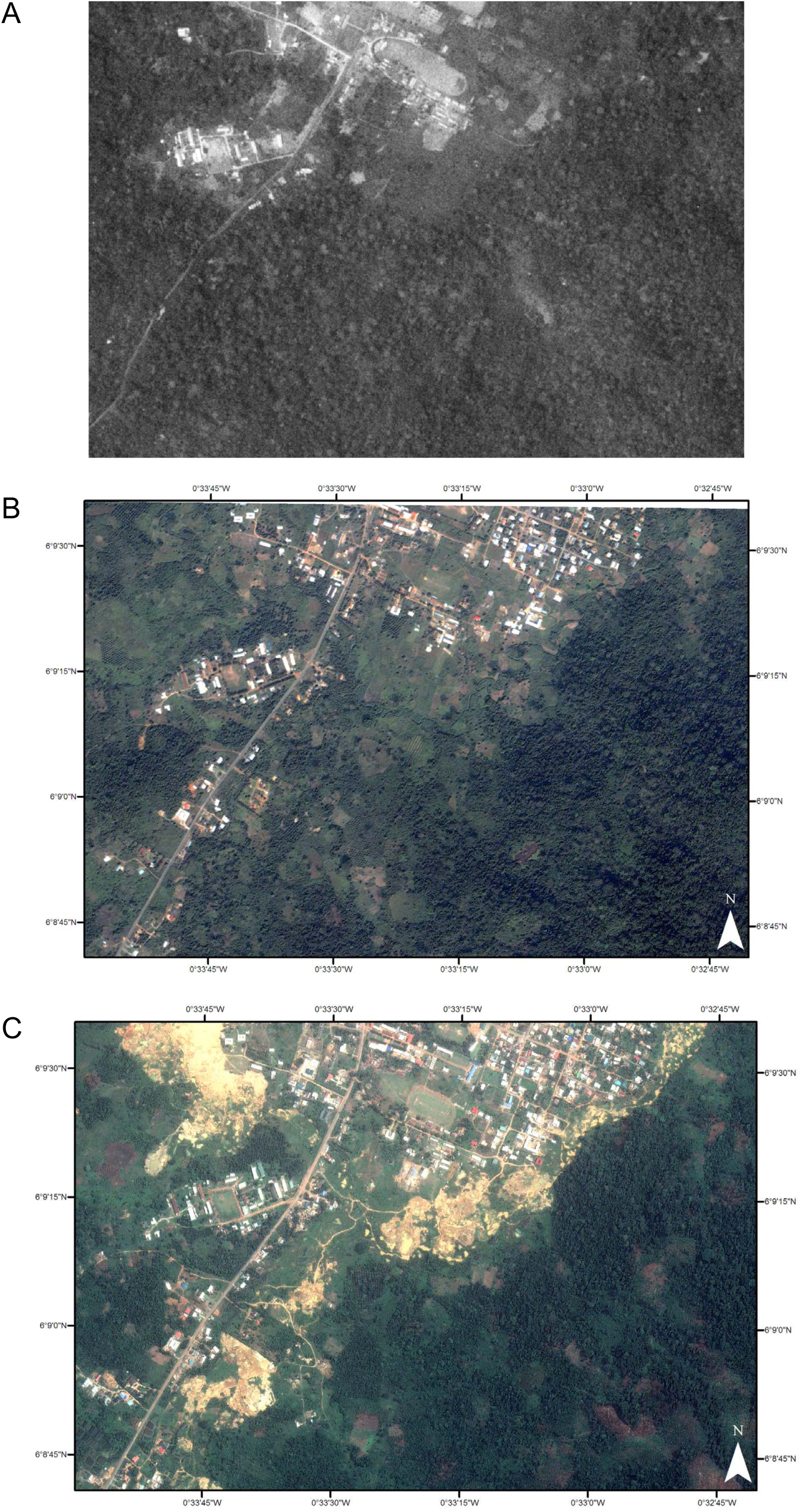
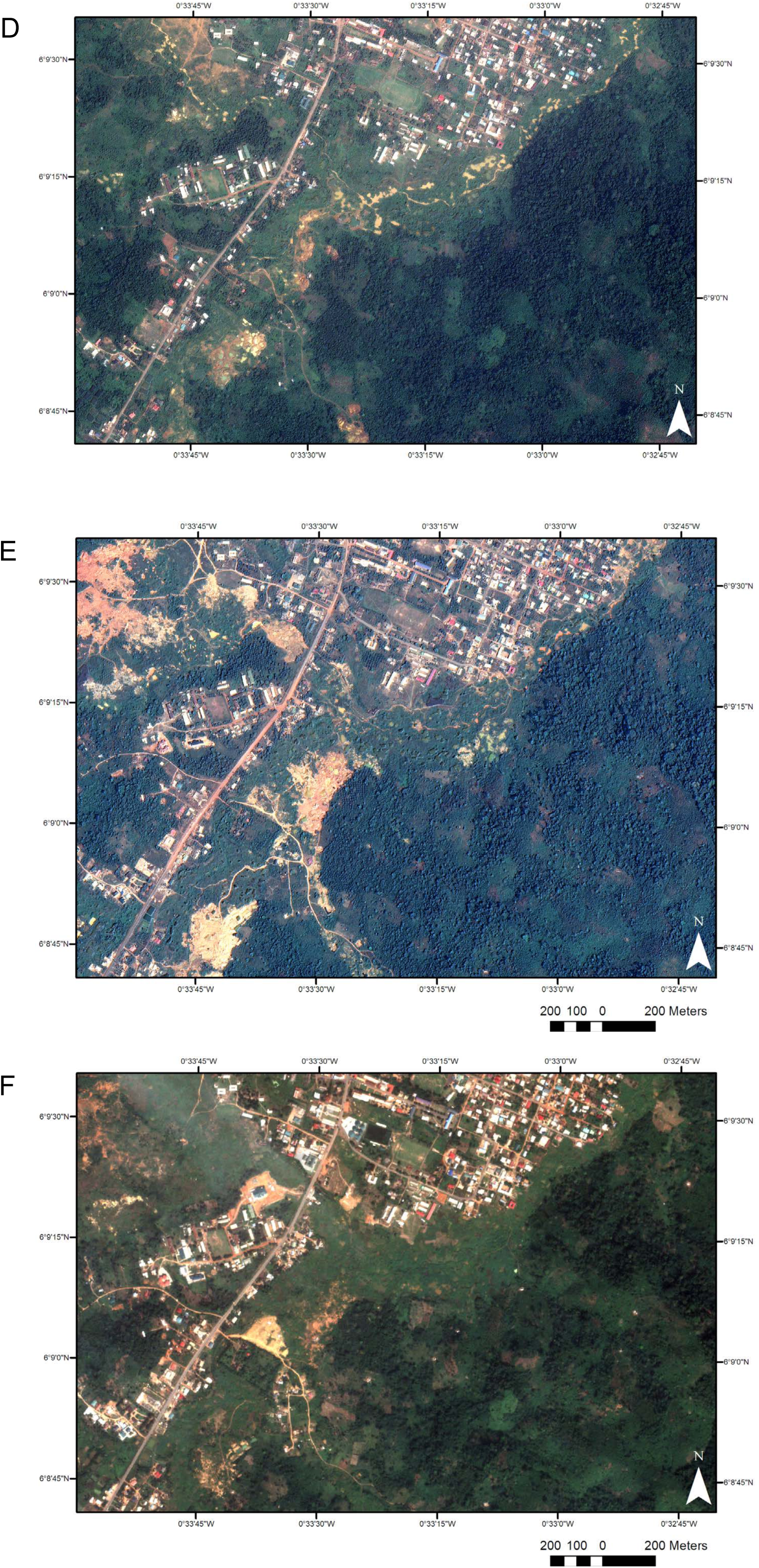
A) Declassified 70 mm film satellite reconnaissance photograph from Jan 28, 1968; B) Pansharpened IKONOS image from Jan 4, 2008; C) Pansharpened Pleides 1B image from March 3, 2014; D) Pansharpened Pleides 1A image from Dec 20, 2014; E) Pansharpened Pleides 1B image from Jan 26, 2017; F) Pansharpened Pleides 1B image from Dec 9, 2018.

**TABLE 1.**
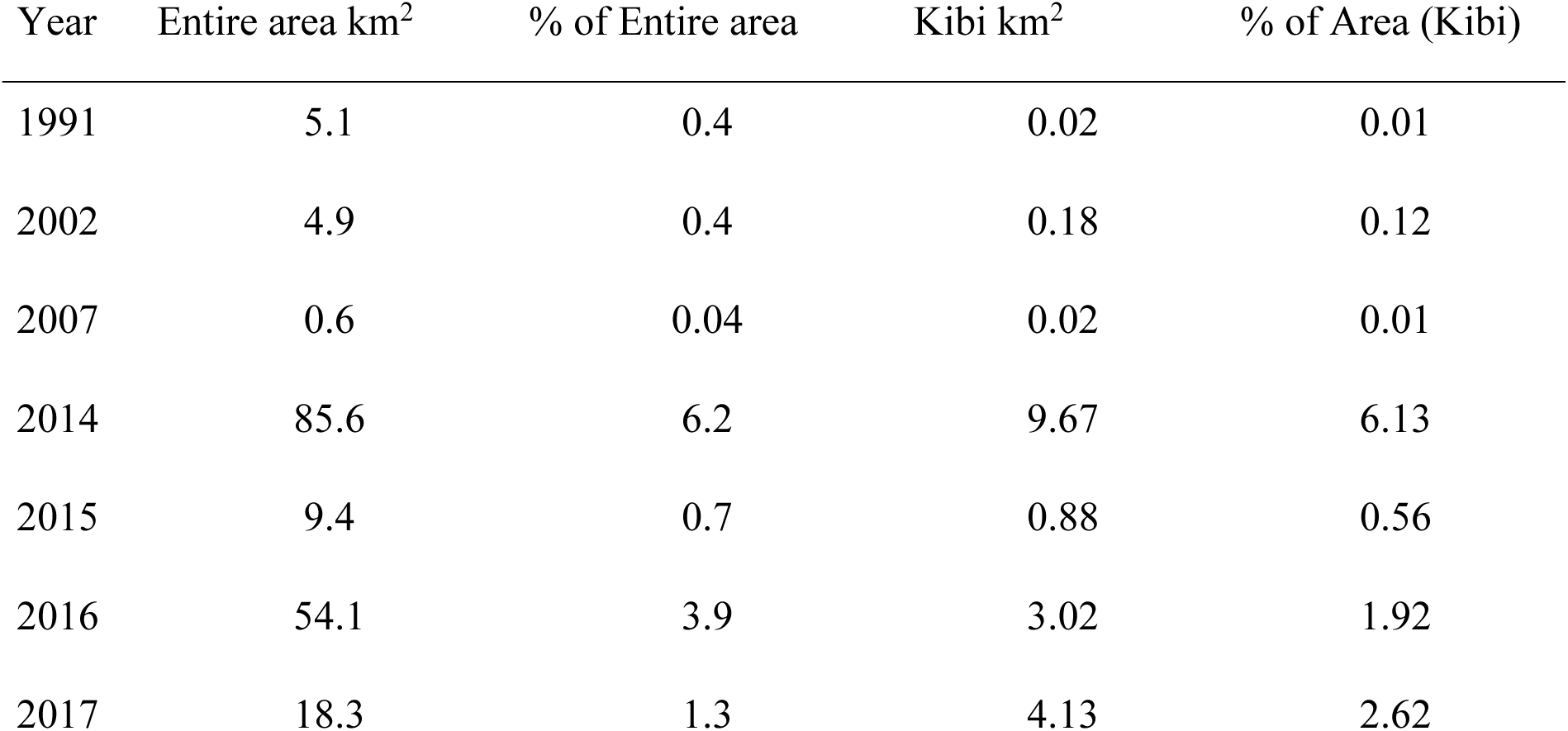
Total area deforested by year for the entire study area (1389.5 km^2^) and around Kibi (157.7 km^2^). Values are reported both in km^2^ and by percent of the area.

**TABLE 2.**
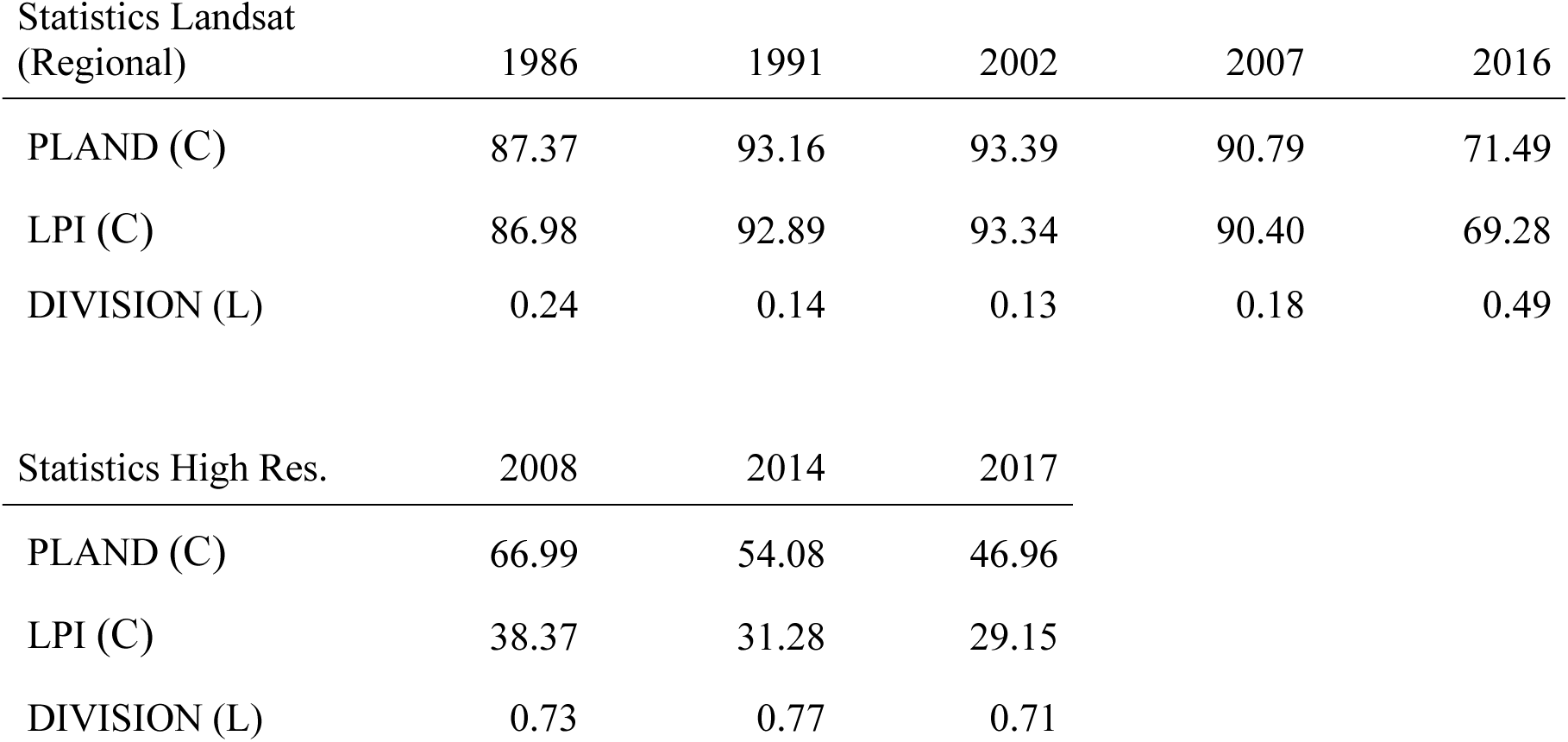
Class and landscape fragmentation statistics for the forest class between 1986 and 2016 for Landsat and from 1991 to 2017 from high resolution imagery (IKONOS and Pleides). PLAND = percentage of landscape, LPI = largest patch index, DIVISION = landscape division index. C denotes class level statistics, L denotes landscape level statistics.

Yearly cloud cover was extracted from the ‘MODIS/Terra Surface Reflectance Daily L2G Global 1 km’ data (Vermote and Wolfe 2015). Average nighttime composite radiance imagery from the ‘Visible Infrared Imaging Radiometer Suite (VIIRS) Day/Night Band (DNB)’ was downloaded from Google Earth Engine for the 25 km^2^ around Kibi for 2012 and 2017 as a proxy for quantifying urban development. Elevation was determined from the Shuttle Radar Topography Mission (SRTM) Digital Elevation Data v4 (Jarvis et al. 2008).

### POPULATION ESTIMATES

We repeatedly explored the study area in 1991, 1993, 2016 and 2018 (February to beginning of April) for population studies of *L. robertsi* as well as for collecting information about the ecological status of the area. In 1991, four people surveyed the creeks surrounding Kyebi up to Asiakwa over a period of six days from their confluence with the Birim River to the higher elevation steep cascades. In 1993, 23 people surveyed the same area for eight days. In 2016, three people surveyed for four days and in 2018, four people surveyed the same area for three days with an additional day in spent in the Ankasa region. Species abundance was estimated through combination of direct visual observation and dip netting. In 1993 electrofishing was also employed. Along the Black and Red Krensen creeks (all four years) as well as creeks along the road to Asiakwa (1991, 1993) higher elevation areas (> 380 m) above the steep cascades were also surveyed. As described by Helfman (1983) and Lucas and Barras (2000), the main benefit of visual observation as a capture independent method of studying fishes is that as long as disturbance is minimized, fish generally behave normally. In the shallow, clear streams, specimens could be readily seen. During the surveys in February, water conductivity was measured several times a day with a Bischof conductivity meter.

### CLASSIFICATION

The Landsat images were classified into forest, non-forest and masked (cloud and water) classes with CLASlite v3.2 (Asner et al. 2009) (Fig. S5A). The main aspect of CLASlite is referred to as Automated Monte Carlo Unmixing (AutoMCU) through which the proportional contribution of photosynthetic vegetation (PV), non-photosynthetic vegetation (NPV) (e.g. branches) and bare substrate (S) to each pixel is determined (Asner et al. 2009). The output is a fractional cover band of the proportions of PV, NPV and S in each pixel. Classification is based on a decision tree of the fractional cover band of the three components. For forest, PV ≥ 80 AND S < 20 while for the non-forest class, PV < 80 OR S ≥ 20. A proportion of 100 would indicate the entire pixel is composed of that component. Forest cover change considered the initial deforestation event (i.e. first year of change).

For the most recent classification (2017) we generated 500 random points throughout the 1389.5 km^2^ study area to assess the thematic accuracy through visual interpretation of a combination of high-resolution imagery available in Google Earth (entire study area) and the Pleides 1B image (Table S2, Fig. 4E). Because the study area is relatively small (< 500,000 ha) and there are only two classes of interest (forest and non-forest), for the classifications from previous years, we generated 150 random points per year and assessed the accuracy through visual interpretation of the imagery (Congalton 2015). Each point represented a homogeneous cluster of at least 3 x 3 pixels. We calculated the overall accuracy (proportion of correctly classified reference points), producer’s accuracy (PA), user’s accuracy (UA) and F-score metrics for each classification (Congalton and Green 2009; Xiong et al. 2017). The PA represents the proportion of reference data that is correctly classified (complement of the omission error). The UA represents the reliability of the classification (complement of the commission error). The F-score measures the accuracy of a class using precision and recall measures (Xiong et al. 2017).

To determine fine scale forest cover change, we carried out a Geographic Object Based Image Analysis (GEOBIA) on the high spatial resolution satellite imagery (Table S2). The objective of GEOBIA is to improve and replicate human interpretation of the imagery in an automated manner (Johansen et al. 2014). GEOBIA has been shown to be highly effective for land cover classification of high spatial resolution imagery (e.g. < 5 m) with low spectral resolution (e.g. 4 – 6 bands) (Chen et al. 2018; Ma et al. 2017). The images were first atmospherically corrected in ENVI 5.5 with FLAASH (Kaufman et al. 1997; Matthew et al. 2000). The imagery from 2008, 2014 and 2017 (March) were then classified with eCognition Developer v9.4 (Trimble Geospatial, Sunnyvale, CA) (Fig. S5B). The first step in the GEOBIA is segmentation. We used parameter of scale = 100, shape = 0.3 and compactness = 0.8. Segmentation groups pixels close in space that are more similar to each other than to pixels in neighbor segments. Next, training samples representing ‘forest’, ‘non-forest vegetation’(NFV), and ‘other’ were manually selected. The class ‘other’ included exposed soil, water and the built-up areas. The images from Dec 2017 and 2018 were not included due to extensive cloud cover in the south-east corner of the images. Object based metrics including the mean surface reflectance in all 4 bands, brightness and texture were calculated for each of the segments. These metrics served as the training data for a nearest neighbor classification. Next, a set of standard classification refinement rules were implemented (Fig. S5B). Fifty forest and 50 non-forest validation points were generated for each year. These were visually assessed for thematic accuracy from the imagery.

### FRAGMENTATION

A fragmentation analysis was performed using Fragstats v4.2.1 (McGarigal et al. 2012) to assess temporal forest fragmentation trends. Class area (CA) and percentage of landscape (PLAND) are important measures of landscape composition; here, they are an indication of how much of the landscape is comprised of forest. In this study, CA is a good indicator of the total forest area lost over time, while PLAND allows for comparison between years of the percent of forest lost over time. The Largest Patch Index (LPI) and the Landscape Division Index (DIVISION) were also calculated. LPI measures the change in the largest forest patch, while DIVISION is a standardized measure of the probability of two randomly selected pixels in the landscape not being located in the same patch of the corresponding patch type (McGarigal et al. 2012). DIVISION ranges 0 - 1, with a value of 0 when the landscape is comprised of a single patch. These metrics were calculated for a 157.7 km^2^ cloud free area around Kibi from the Landsat classification for 1986, 1991, 2002, 2007 and 2016. They were also calculated for the high-resolution imagery classified with the GEOBIA approach.

Foreground Area Density (FAD), a spatial fragmentation measure, was calculated in 20 m buffers from high spatial resolution classifications around the three Krensen creeks with the GUIDOS Toolbox (Vogt and Riitters 2017). Twenty meters is the upper limit of the recommended buffer zone width in Ghana for minor perennial streams (Ministry of Water Resources Works and Housing 2013). FAD measures the proportion of forest pixels at five observation scales from 7 to 243 pixels. Results are classified into six classes: < 10% (rare), 10-39% (patchy), 40-59% (transitional), 60-89% (dominant), 90-99% (interior), 100% (intact). The proportion of the six FAD classes was calculated for the buffers as the average from the five observation scales (Vogt and Riitters 2017).

## RESULTS

In comparison to the baseline year of 1986, the forest cover change analysis for the entire study area (Table 1) shows the greatest increase in initial deforestation events in 2014 and 2016 (6.2% and 3.9% of the area respectively). Deforestation events also increased in 2017 (1.3% of the area). The overall spatial pattern indicates an easterly progression of the deforestation over time (Fig. 5). In 2014, the deforested areas were not only concentrated around the river and creek banks but also in patches west of the Atewa Forest Reserve. In 2016, disconnected patches of deforestation can be seen in the eastern sector of the study area. The steep western slopes of the Atewa range remained unaffected with encroachment seen on the eastern slopes beginning in 2014 and progressing to higher elevations in 2016 and 2017. Around Kibi, the main deforestation events in 2014 (6.13% of the area) follow the river and creeks with disconnected patches of forest clearing in 2016 and 2017 (1.9 and 2.6% of the area respectively) (Table 1). The classifications for all years had a high level of accuracy for both forest and non-forest classes ranging from 82 percent to 98.8 percent (Table S3). The lowest accuracy was for the PA of the non-forest class from the 2014 image (82%, 18% omission error). The highest accuracy was for the forest class from the 1986 image (PA = 98.8%, 1.2% omission error). The F-score ranged from 0.84 to 0.96. The GEOBIA also resulted in high classification accuracies ranging from 86 percent to 100 percent (Table S4). The lowest accuracy was for the UA of the forest class from 2017 (86%, 14% omission error). The highest accuracy was for the PA of the forest class from 2017 (PA = 100%). The F-score ranged from 0.92 to 0.94.

**FIGURE 5.**
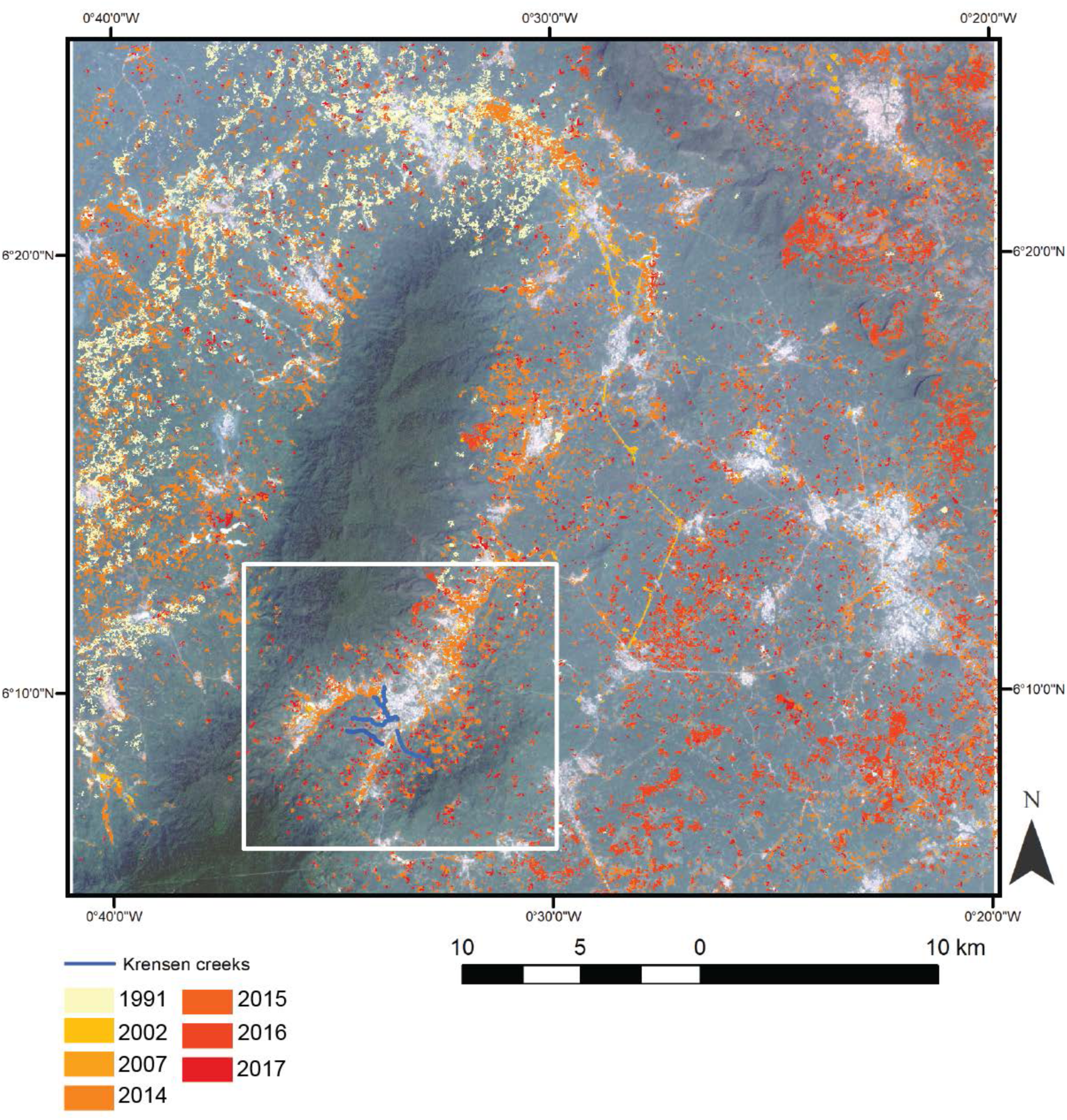
Deforestation by year of observation in the 1389.5 km^2^ area of the Birim river basin around the Atewa forest reserve as determined from the Landsat imagery. The patches of deforestation can be seen to have an easterly progression. Extensive deforestation is seen at low elevations around Atewa Forest reserve along the banks of the Birim river and small creeks in 2014 indicative of AGSM activities. Patchy deforestation indicative of agricultural activities can be seen in the eastern lowlands from 2016. Deforestation is seen at higher elevations along the slopes of the Atewa Forest reserve and east of Kibi in 2017.

**TABLE 3.** Summary of aquatic habitat fragmentation agents acting upon the Krensen creeks based on Fuller et al. (2015). Agents with an * are a result of Riparian forest clearing. The specific type of ‘waterfall’ present in the study area is the ‘steep cascade’.

| **Fragmentation agent** | **Origin** | **Barrier form** | **Effect on *L. robertsi*** |
| --- | --- | --- | --- |
| Waterfall | Natural | Physical | Impassable barrier |
| Predatory/Competitive species | Natural/<br>anthropogenic | Biological | Predation of individuals.<br>Competition for resources and habitat. |
| Pollution (agriculture and AGSM)* | Anthropogenic | Habitat | Decrease in water quality leading to decrease in available habitat. |
| Increased nighttime lights* | Anthropogenic | Habitat | Negative impact on vision. |
| Increased water temperature* | Anthropogenic | Habitat | Negative impact on sex ratio. |

Table 2 and Fig. 6 reveal the forest fragmentation patterns that the area has experienced. Based on the Landsat imagery, for the 157.7 km^2^ around Kibi, the largest decrease in forest area is seen in the last period between 2007 to 2016 during which there is also an increase in the number of forest fragments from 160 to 445 (Fig. 6). From 1986 to 2007 forest cover was relatively stable, including marginal forest gain between 1986 to 1991 (PLAND from 87.4% to 93.2%), negligible change from 1991 to 2002, and a small decrease in forest area between 2002 and 2007 (PLAND from 93.4% to 90.4%). There was however, a large increase in the number of fragments between 2002 and 2007 (41 to 160). Similar trends are shown by the LPI metric, which revealed a considerable reduction of the largest patch from 90.4 percent in 2007 to 69.28 percent in 2016 (Table 2). At the landscape level, the DIVISION metric indicates an increase in 2016 (0.49) compared to previous years (e.g. 0.24 for 1986). In addition, we found that the deforestation progressed to higher elevations over time. In 1986, 50 percent of the non-forest class was located at elevations below 319 m (μ = 333 m ± 65 m) with 95 percent below 460 m. In 2016, 50 percent of the non-forest class had increased in elevation to areas below 335 m (μ = 355 m ± 83 m) with 95 percent below 517 m.

**FIGURE 6.**
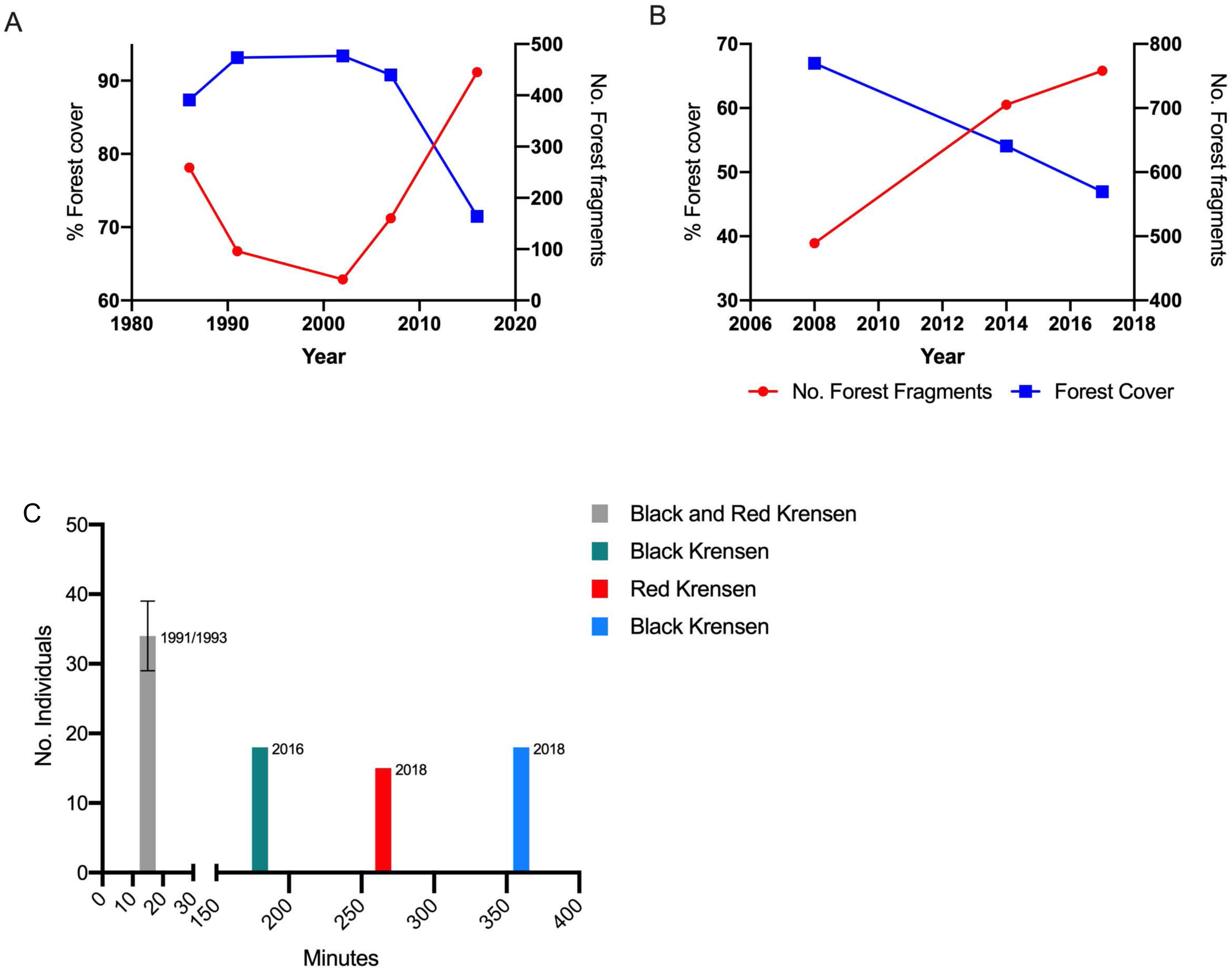
A) Landsat image classification from 1986 to 2016 indicating total forest area and number of forest fragments within the 1389.5 km^2^ area; B) High spatial resolution IKONOS (2008) and Pleides (2014 and 2017) image classification indicating total forest area and number of forest fragments within the 157.7 km^2^ around Kibi; C) Time (min) needed to observe a minimum of 15 *L. robertsi* in the 1990s compared to 2016 and 2018 in the Black and Red Krensen creeks. In 2016 and 2018 *L. robertsi* was only seen at elevations higher than 360 m.

The forest loss trend from the high-resolution GEOBIA (Fig. 6, Table 2) is similar to that of the landscape scale. For instance, there is a decrease in forest area from 16.23 km^2^ (2008) to 11.60 km^2^ in 2017 (PLAND from 67.0% to 47.0%). The number of patches also increases significantly from 227 patches in 2008 to 705 patches in 2014 and 758 in 2017 (Fig. 6). The LPI reveals a reduction of the largest forest fragment for the same three periods (35.0%, 31.3% and 29.2%, respectively) (Table 2).

For the 25 km^2^ around Kibi, the nighttime satellite imagery from the VIIRS DNB data indicates the artificial illumination from settlements more than doubled from 2012 μ=0.29 ± 0.75 nW/cm^2^/sr (range: 0-6.01 nW/cm^2^/sr) to 2017 μ=0.71 ± 0.89 nW/cm^2^/sr (range: 0.13-6.73 nW/cm^2^/sr). At a finer grain, along the elevation gradient of the Black Krensen, the greatest increase in nighttime illumination is seen at elevations > 360m: in 2012, 0.07 – 0.09 nW/cm^2^/sr were observed increasing to 0.4-0.5 nW/cm^2^/sr in 2017. For comparison, the centre of Kyebi increased from 5.78 to 6.73 nW/cm^2^/sr over the same time period.

The length of time needed to record the presence of individuals increased greatly over time (Fig. 6C). In the 1990s, over 30 individuals were recorded within 15 min of sampling. In 2016, 3 hrs were needed to record 18 specimens in the Black Krensen. The time further increased to 6 hrs in 2018. From the combined field observations in 2016 and 2018 we estimated the Kibi area *L. robertsi* population size to be less than 1000 individuals spread over the three small Krensen creeks which are less than 2 m wide and 0.5 m deep. Compared to our observations from the 1990s, this represents approximately ten percent of the original population. As seen in the field photographs from 2016 and 2018 the quality of the habitat around the creeks has decreased considerably compared to 1991 (Fig. 2). From our measurements, conductivity in the Black Krensen increased from 220 µS/cm (1991) to 288 µS/cm (2018). Wastewater, urban runoff, agricultural runoff, soil erosion and tailings are major sources of pollution which can increase conductivity. No individuals of *L. robertsi* were found in the Ankasa region in 2018 and as such the persistence of the species there could not be confirmed.

No cichlids were found above the steep cascades, only *Enteromius walkeri* (barb), *Epiplatys chaperi* (killifish) and *Amphilius atesuensis* (catfish). Similarly, northwest of the type locality (Fig. S6, S7) within the buffer zone of the Atewa Forest Reserve, the water in the creeks remains clear but *L. robertsi* was not found due to the steep cascades (Fig. S6) which the cichlid cannot cross. Based on the description of the species, in 1965 when the type specimens were collected, the habitat consisted of a series of shallow pools with relatively quiet water connected by riffles with a lack of algae or vascular aquatic plants. The banks, overgrown with broadleaved herbaceous plants (up to one m tall) provided shade to the stream (van den Audenaerde and Loiselle 1971). Similar conditions can be seen in the field photograph from 1991(Fig. 2). Most of the creeks along the road between Kibi and the type locality west of Asiakwa (Fig. 1B) no longer existed in 2016, they had been destroyed by new road development or ASM operations (Fig. S6).

In 2016 and 2018 no *L. robertsi* were observed in the lower portions of the Krensen creeks near their confluence with the Birim river. Here, in 2016 *Coptodon* sp. was found, whereas it had not been present in the 1990s. It has a higher reproduction rate as a free substratum spawner than the cave/mouth brooding *L. robertsi*. In 2018, we also observed *Hemichromis fasciatus* and *Parachanna obscura* in the lower portions of the creeks. In 1991 and 1993 *L. robertsi* was present from approximately 320 m elevation and higher, while in 2016 and 2018 the lowest elevation it was observed was 360 m. These elevation values are consistent with the increase in elevation observed in from the deforestation and FAD analyses. In the 320-360 m altitude zone where *L. robertsi* was no longer observed in 2016 and 2018, the FAD classes within the creek buffer zones indicate an increase in non-forest proportion from 59.7% in 2008 to 64.6% (2014) and 77.5% (2017) (Figure 7A). A large decrease in the dominant class can also be seen from 22.7% (1991) to 17.3% (2014) and 11.8% (2017). Applying a minimum of 15% canopy cover from Ghana’s forest definition (National REDD+ Secretariat Forest Commission 2017), forest decreases within the buffer zones in the 320-360 m altitude range from 40.2% (2008) to 35.4% (2014) and 22.5% (2017). Combined aquatic habitat fragmentation agents (Table 3) are shown in Figure 7B for the Black Krensen. At the highest elevation an impassable natural barrier consists of steep cascades. The extent of *L. robertsi’*s distribution has retreated upstream, reduced by anthropogenic habitat barriers at low elevations where the Riparian buffer is deforested and there is pollution form ASGM activities and agricultural runoff. Increased development illustrated by the illumination from nighttime lights further poses a habitat barrier to the species. Predatory/competitive species form a recent biological barrier at lower elevations (not present in the 1990s). As a result, the core creek length with suitable habitat for *L. robertsi* has decreased with the lower limit observed to be 360 m elevation in 2018. The estimated loss of core habitat length from the 1990s to 2018 due to the natural, anthropogenic and biological fragmentation agents is 950 – 980 m (Fig. 7B,C).

**FIGURE 7.**
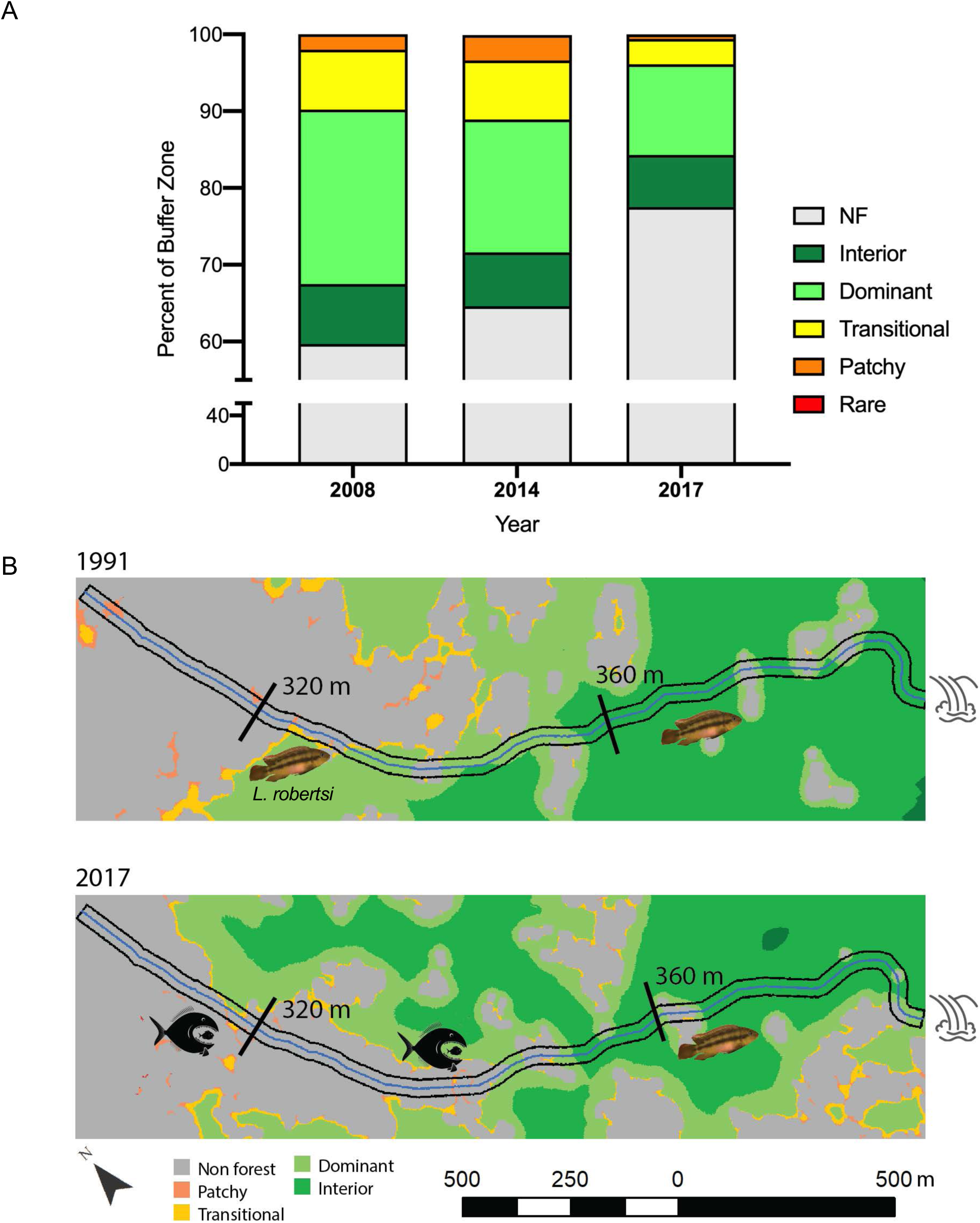
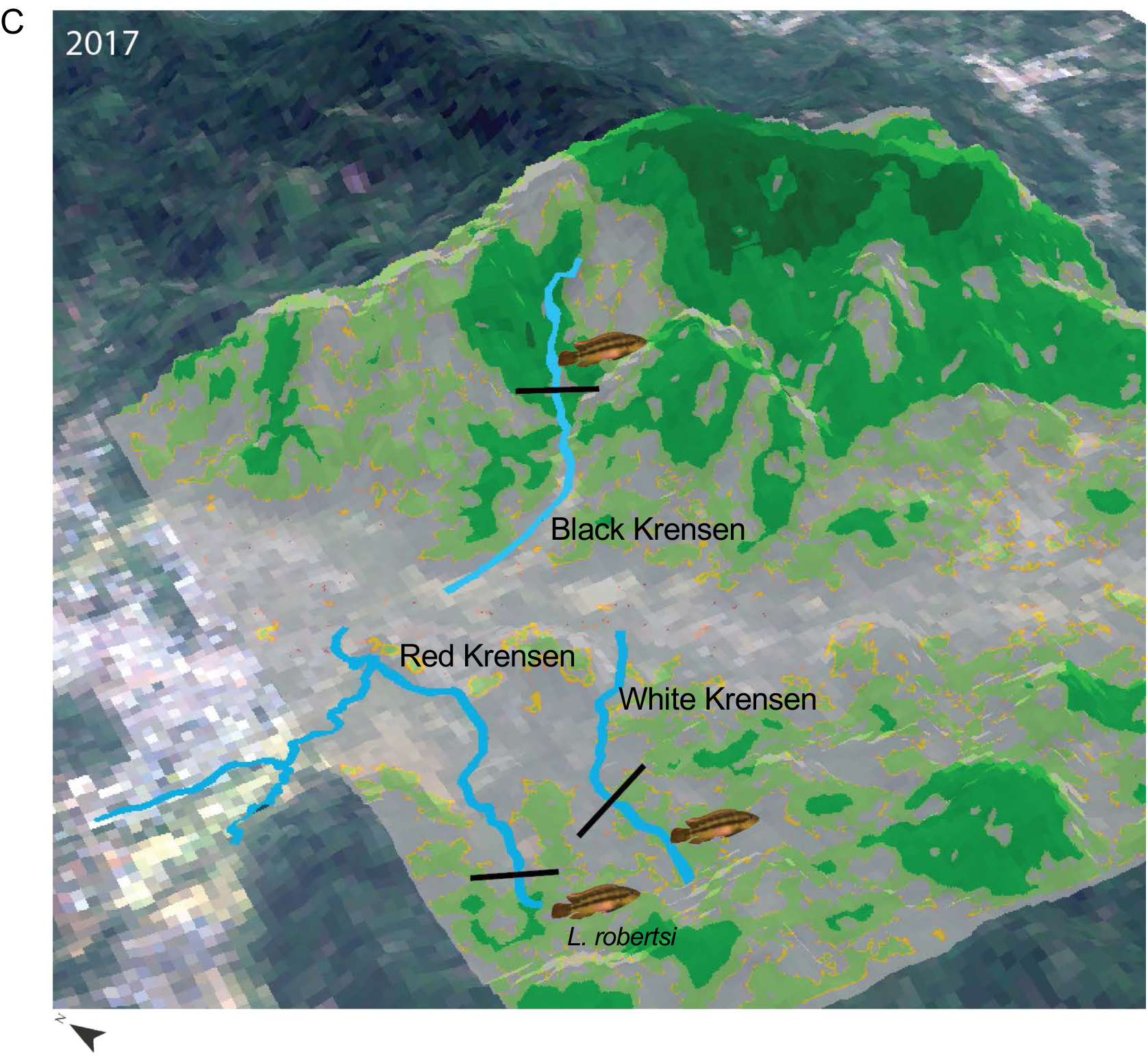
A) Change in proportion of Foreground area Density (FAD) forest fragmentation classes in 20 m buffers around the Black, Red and White Krensen creeks. The values represent the average of 5 observation scales (window sizes ranging from 7 to 243 pixels). The classes represent the amount of forest within the average window size as < 10% (rare), 10-39% (patchy), 40-59% (transitional), 60-89% (dominant), 90-99% (interior). The sixth class of 100% (intact) was not encountered. These results are based on the high spatial resolution forest classification. B) Aquatic habitat fragmentation agents affecting the *L. robertsi* habitat in the Black Krensen (with 20 m buffer) is illustrated. These include physical barriers (steep cascades) at high elevations, decreased water quality due to Riparin forest loss from AGSM and agricultural activities and the emergence of predatory/competitive species at low elevations. The 320 m and 360 m elevation thresholds indicate the lowest limit where *L. robertsi* was found in the 1990s and 2016/18 respectively. C) FAD classes from 2017 superimposed upon Landsat image and a digital elevation model. The Krensen creeks with the 360 m elevation limit are shown below which *L. robertsi* is no longer found. The Riparian forests along the creek banks are the most intact in areas with the steepest slopes.

## DISCUSSION

### HABITAT LOSS AND FRAGMENTATION

While currently classified as *endangered* by the IUCN Red List, due to the limited range and very small number of specimens left in the wild (Entsua-Mensah and Laleye 2010), *L. robertsi* is in highest need for a conservation program and should be considered as close to extinction with its IUCN Red List classification upgraded to *critically endangered*. As our results show, its habitat has undergone substantial degradation since the last IUCN assessment primarily from ASGM activities (Schep et al. 2016). Based on our most recent field observations, *L. robertsi* is absent in the lower parts of the creeks (< 320 m elevation) (Fig 7B,C), and the Birim River. While we observed *L. robertsi* occurring in these lower parts of the creeks in small numbers in the 1990s, due to the changes in the aquatic environment including both anthropogenic and biological habitat fragmentation agents (Fig. 7B,C, Table 3), none were found in these areas in 2016 or 2018. Instead, we observed the presence of highly competitive and/or predatory species (e.g. *H. fasciatus*, *P. obscura*, *Coptodon* sp.) which can readily outcompete *L. robertsi*. Overall, the range of the creeks where *L. robertsi* occurs today has decreased by 950 – 980 m in length (representing less than 50% of its extent in the 1990s). The FAD results indicate that within the 20 m buffer around the creeks in the 320-360 m elevation range where *L. robertsi* is no longer present, deforestation has increased to nearly 80%. Interior forest (90-99% forest cover) surrounds only 7.8% of the creeks in this elevation range (Fig. 7A). In addition to the low elevation predatory/competitive species, increased water temperature caused by the low proportion of Riparian tree cover, decrease in water quality from sedimentation, runoff and ASGM activities and increased light pollution are all likely the factors responsible for *L. robertsi* becoming confined to higher elevations of the creeks. The decline in Riparian forest extent along the buffer zones of the creeks has further led to habitat fragmentation resulting in less exchange of genotypes between individuals within the same habitat and lesser availability of territories for breeding pairs (*L. robersti* needs a larger territory for breeding than *Coptodon* sp). The shorter stretches of the creeks where they are now found is reducing the space for breeding pairs.

### FOREST COVER CHANGE

Our results demonstrate an accelerated loss of forest cover over the 1986-2017 period with the largest initial deforestation events visible in 2014 and 2016 (Fig. 5,6). These results are based on the first observation of deforestation. Due to persistent cloud cover in the area, suitable imagery was not available in additional years. On average, there were fewer than 60 clear days per year (Fig. S8). Therefore, some of the deforestation recorded in 2014 may have occurred between 2008 and 2013. This accelerated degradation of the landscape has also been reported by Meijer et al. (2018) and Tappan et al. (2016) with attribution to agricultural expansion and mining. For the area around Kibi, deforestation for agriculture can be seen in the high-resolution imagery from 2008 (Fig. 4B) in comparison to the satellite photograph (1968) which shows intact forest (Fig. 4A). Nevertheless, in 2008, Riparian vegetation remained along the banks of the river and creeks. In contrast with the imagery in 2014 and 2017, the Pleides image from December 2018 (Fig. 4F) illustrates an abandonment of the majority of the ASGM sites and an emergence of low stature herbaceous vegetation (Fig. 2F). Similarly, the type locality (approx. 15 km north of Kibi) which had been forested in the 1960s is an abandoned ASGM pit with regenerating herbaceous vegetation (Fig. S7).

Forest gain and loss in tropical areas can be shaped by internal and external socio-economics forces that result in variations in forest cover which can be described by a forest fragmentation analysis (e.g. Arroyo-Mora et al. 2005). At the landscape level our fragmentation analysis (Table 2) indicated an increase in metrics PLAND, and LPI along with a simultaneous decrease in landscape division from 1986 to 1991 potentially related to old secondary forest growth in abandoned fields (e.g. 17 to 20 year old-growth). It could also be to some degree an artifact given the differences in sensor characteristics between the older generation Landsat TM4 and Landsat TM5 sensors (Micijevic et al. 2016). Scale is an important factor in landscape ecology directly influencing the results and interpretation of the fragmentation statistics. Fragmentation patterns (Fig. 6A,B) for the study area at two spatial scales are consistent and show an increase in the number of patches, and a reduction in PLAND and LPI with a simultaneous increase in landscape division up to 2002, with a greater change of these metrics from 2007 to 2016. Such changes in the landscape structure are typical for areas with progressive fragmentation due to LUCC practices related to the expansion of the agricultural frontier (e.g. de Oliveira et al. 2017; Su et al. 2014), road network and/or urban development (Liu et al. 2013).

Tropical forest classification with CLASLite is well established and studies have repeatedly shown high accuracies (e.g. (Allnutt et al. 2013; Arjasakusuma et al. 2018; Asner et al. 2013; Chicas et al. 2016; da Silva et al. 2018)). Likewise, we obtained high thematic map accuracies for our eight classifications (Table S3). A challenge with determining the accuracies of our classifications was the temporal component (31 years). No pre-existing reference data were available from other sources for the years used in this study. However, with CLASLite, the training of the classification was independent from the collection of the reference data used to evaluate accuracy (Congalton 2015). Classification is based solely on a decision tree executed on the fractional components of PV, NPV and S derived from the AutoMCU process (Asner et al. 2009). None of the misclassified pixels (Table S3) were in areas with exposed sediment or water with high sediment load (e.g. tailings ponds and soil from ASGM), they were all in areas with degraded vegetation, early regrowth, mixed agriculture, trees crops, agroforestry, or oil palm plantations where the differentiation of forest versus non forest based on the decision tree rule was difficult to determine for the visual interpreter. In addition, differences in sensor characteristics also contribute to variable image quality between the Landsat satellites over the years (Markham et al. 2018).

### SOCIO-ECONOMIC DRIVERS

The World Bank estimates Ghana’s economy expanded by 8.5 percent in 2017 driven primarily by the mining and oil sectors. Gold mining around the Atewa Forest Reserve accelerated as of 2009 (RMSC 2016) coinciding with a sharp rise in the value of gold to a high of $1514/troy oz in 2012 (World Bank 2018) following the 2008 global economic recession. Furthermore, the temporary Ghanaian diamond export ban led to a large number of miners switching to ASGM by 2007 (Nyame and Grant 2012). From the agricultural sector, an increase in cocoa production has been responsible for large scale forest clearing as the traditional shade dependent variety is replaced by open field hybrid varieties which require full sun (Kolavalli and Vigneri 2011). In the Eastern region of Ghana, 50 percent of the cocoa grown is the full sun variety (Kolavalli and Vigneri 2011; Obiri et al. 2007). International cocoa prices were highest in 2015 (3.2$/kg) (World Bank 2018). The deforestation events which expanded in 2016 and 2017 in discrete plots away from the rivers and creeks (Fig. 5) may be related to increases in local consumption as well as international agricultural commodity prices.

Recently, the Government of Ghana negotiated an agreement with Sinohydro Corporation of China whereby Sinohydro will provide $2 billion worth of infrastructure, in exchange for proceeds of refined bauxite (Government of Ghana 2018, International Monetary Fund 2019). Despite a recommendation from the IUCN and Ghanaian and international conservation organizations to increase the status of the Atewa Forest Reserve to a National Park as a way to maximize sustainable ecosystem services, the Atewa Forest Reserve is part of the infrastructure for natural resource exchange; the mountain range is estimated to contain 200 million tons of bauxite (Meijer et al. 2018; Schep et al. 2016).

### CONCLUDING REMARKS

We have shown the potential for multi-scale, multi-temporal remote sensing derived ecosystem attributes to inform processes controlling the distribution of a restricted range cichlid in Ghana. Ecosystem attributes are well known robust predictors of biodiversity change (Arenas-Castro et al. 2019). Efforts in mapping Essential Biodiveristy Variables (EBVs) from satellite imagery have identified both primary and secondary/proxy variables that can be either directly mapped or inferred from remotely sensed data (Vihervaara et al. 2017, Kissling et al. 2018). Similarly, we map changes in natural and anthropogenic fragmentation agents (Fuller et al. 2015) acting upon the reaches of creek habitat important for *L. robertsi* (Fig. 7). Field visits confirm a loss in overall abundance of the species with an increase in predatory and competitive species. The spatio-temporal nature of the multi-resolution satellite imagery (Fig. 4,5) further allowed for inferring socio-economic drivers of the LUCC and habitat fragmentation.

While *in-situ* conservation of the habitat would be best, extensive remediation of the remaining habitat around Kibi and the Ankasa region (if the species still exists there) would be needed. Further exploration of both areas, including higher elevation creeks within the Atewa Forest Reserve, is needed to confirm exact distribution and population size. If the planned extraction of bauxite (large open pit mine) proceeds, most of the potential habitat could be lost in a very short time. An *ex-situ* program, in coordination with scientific institutions out of Ghana (e.g. Ministry of Fisheries and Aquaculture Development), as well as public aquariums, zoos and private specialists seems to be the most effective way to stop the extinction of this Ghanaian endemic. Such a program has been started in Austria with participation from the Vienna Zoo and other institutions. Also, a campaign to inform those living in close proximity to the Krensen creeks, that a species found nowhere else on the planet is in their backyard may increase the chances to protect *L. robertsi* and its native habitat from further damage.

## ACKNOWLEDGEMENTS

We would like to thank Edward Gyekety (Regional Cartographer, Forest Services Division, Koforidua, Eastern Region), Ransford Tetteh (Surveyor, Koforidua), and Adjei Joseph (Koforidua) (Ghana) for collecting field photographs in 2018 and 2019, Patrick Boateng (Kibi) for field assistance as well as Paul Loiselle (USA) for information about the habitats around Kibi in the 1960s, Ulrich Schliewen (Germany) for information from the Ankasa region (1990s) and Adrian Indermauer (Switzerland) for the latest population update in 2018. We also thank Eddie K. Abban (Ghana) for his help in 1991 and 1993, Andreas Wanninger (Vienna) for support of the work of A. Lamboj, and Anton Oberleuthner (Austria) and Manuel Zapater (Spain) for their help with observations in the field. Remote sensing analysis was supported by the Natural Sciences and Engineering Research Council of Canada.

## DATA AVAILABILITY

The Landsat satellite imagery used in this study can be downloaded at no cost from the USGS Earth Explorer: https://earthexplorer.usgs.gov/ Similarly, the VIIRS DNB imagery (https://developers.google.com/earth-engine/datasets/catalog/NOAA_VIIRS_DNB_MONTHLY_V1_VCMCFG), MODIS imagery (https://developers.google.com/earth-engine/datasets/catalog/MODIS_006_MOD09GA) and the SRTM Digital Elevation data v4 (https://developers.google.com/earth-engine/datasets/catalog/CGIAR_SRTM90_V4) are available at no cost through Google Earth Engine.

