## Supplementary Material for "Habitat loss in the restricted range of the endemic Ghanaian cichlid *Limbochromis robertsi*"

TABLE S1. Landsat imagery used to assess forest cover change in a 1389.5 $km^2$ area around Kibi. Images were chosen to minimize cloud cover over the study area. Only the 30 m multispectral bands were used in the classification.

| Date (YYYY/MM/DD) | Sensor |
|---|---|
| 1986/12/22 | TM 5 |
| 1991/01/10 | TM 4 |
| 2002/12/26 | ETM+ 7 |
| 2007/01/06 | ETM+ 7 |
| 2014/03/22 | OLI 8 |
| 2015/01/04 | OLI 8 |
| 2016/01/23 | OLI 8 |
| 2017/04/15 | OLI 8 |

TABLE S2. High resolution satellite imagery around the *L. robertsi* collection site south of Kibi. *The same image was also acquired for the type locality.

| Date (YYYY/MM/DD) | Sensor | Spatial resolution for Multispectral (MS) and Panchromatic (Pan) bands | No. bands |
|---|---|---|---|
| 1968/01/28 | 70 mm Panoramic Film | | 1 black and white |
| 2008/01/04 | IKONOS | 3.2 m (MS) / 1 m (Pan) | 4 MS + Pan |
| 2014/03/03 | Pleides 1B | 2 m (MS) / 50 cm (Pan) | 4 MS + Pan |
| 2014/12/20 | Pleides 1A | 2 m (MS) / 50 cm (Pan) | 4 MS + Pan |
| 2017/01/26 | Pleides 1B | 2 m (MS) / 50 cm (Pan) | 4 MS+ Pan |
| 2018/12/09* | Pleides 1B | 2 m (MS) / 50 cm (Pan) | 4 MS+ Pan |

Table S3. Confusion matrices for the Landsat image classifications for a) 1986, b) 1991, c) 2002, d) 2007, e) 2014, f) 2015, g) 2016, h) 2017

A Landsat TM 5 - 1986

| | | Reference | | |
|---|---|---|---|---|
| | | Forest | Non forest | User's Accuracy |
| Classification | Forest | 85 | 8 | 91.4 |
| | Non forest | 1 | 56 | 98.2 |
| | Producer's Accuracy | 98.8 | 87.5 | **Overall 94.0** |
| | F-score | 0.95 | 0.93 | |

B Landsat TM 4 - 1991

| | | Reference | | |
|---|---|---|---|---|
| | | Forest | Non forest | User's Accuracy |
| Classification | Forest | 79 | 8 | 90.8 |
| | Non forest | 9 | 54 | 85.7 |
| | Producer's Accuracy | 89.8 | 87.1 | **Overall 88.7** |
| | F-score | 0.90 | 0.86 | |

C Landsat 7 ETM+ - 2002

| | | Reference | | |
|---|---|---|---|---|
| | | Forest | Non forest | User's Accuracy |
| Classification | Forest | 78 | 8 | 90.7 |
| | Non forest | 6 | 58 | 90.6 |
| | Producer's Accuracy | 92.9 | 87.9 | **Overall 90.7** |
| | F-score | 0.92 | 0.89 | |

D Landsat 7 ETM+ - 2007

| | | Reference | | |
|---|---|---|---|---|
| | | Forest | Non forest | User's Accuracy |
| Classification | Forest | 85 | 4 | 95.5 |
| | Non forest | 4 | 57 | 93.4 |
| | Producer's Accuracy | 95.5 | 93.4 | **Overall 94.7** |
| | F-score | 0.96 | 0.93 | |

E Landsat 8 OLI - 2014

| | | Reference | | |
|---|---|---|---|---|
| | | Forest | Non forest | User's Accuracy |
| Classification | Forest | 81 | 11 | 88.0 |
| | Non forest | 8 | 50 | 86.2 |
| | Producer's Accuracy | 91.0 | 82.0 | **Overall 87.3** |
| | F-score | 0.90 | 0.84 | |

F Landsat 8 OLI - 2015

| | | Reference | | |
|---|---|---|---|---|
| | | Forest | Non forest | User's Accuracy |
| Classification | Forest | 72 | 12 | 85.7 |
| | Non forest | 6 | 60 | 90.9 |
| | Producer's Accuracy | 92.3 | 83.3 | **Overall 88.0** |
| | F-score | 0.89 | 0.87 | |

G Landsat 8 OLI - 2016

| | | Reference | | |
|---|---|---|---|---|
| | | Forest | Non forest | User's Accuracy |
| Classification | Forest | 80 | 6 | 93.0 |
| | Non forest | 7 | 57 | 89.1 |
| | Producer's Accuracy | 92.0 | 90.5 | **Overall 91.3** |
| | F-score | 0.92 | 0.90 | |

H Landsat 8 OLI - 2017

| | | Reference | | |
|---|---|---|---|---|
| | | Forest | Non forest | User's Accuracy |
| Classification | Forest | 335 | 16 | 95.4 |
| | Non forest | 20 | 94 | 82.5 |
| | Producer's Accuracy | 94.4 | 85.5 | **Overall 92.3** |
| | F-score | 0.95 | 0.84 | |

Table S4. Confusion matrices for the high spatial resolution Object Oriented Classifications for a) 1991 b) 2014, c) 2017, d) 2018

A IKONOS - 1991

| | | Reference | | |
|---|---|---|---|---|
| | | Forest | Non forest | User's Accuracy |
| Classification | Forest | 49 | 1 | 98.0 |
| | Non forest | 5 | 45 | 90.0 |
| | Producer's Accuracy | 90.7 | 97.8 | **Overall 94.0** |
| | F-score | 0.94 | 0.94 | |

B Pleides 1B - 2014

| | | Reference | | |
|---|---|---|---|---|
| | | Forest | Non forest | User's Accuracy |
| Classification | Forest | 44 | 6 | 88.0 |
| | Non forest | 2 | 48 | 96.0 |
| | Producer's Accuracy | 95.7 | 88.9 | **Overall 92.0** |
| | F-score | 0.92 | 0.92 | |

C Pleides 1B - 2017

| | | Reference | | |
|---|---|---|---|---|
| | | Forest | Non forest | User's Accuracy |
| Classification | Forest | 43 | 7 | 86.0 |
| | Non forest | 0 | 50 | 100 |
| | Producer's Accuracy | 100 | 87.7 | **Overall 93.0** |
| | F-score | 0.92 | 0.93 | |

A

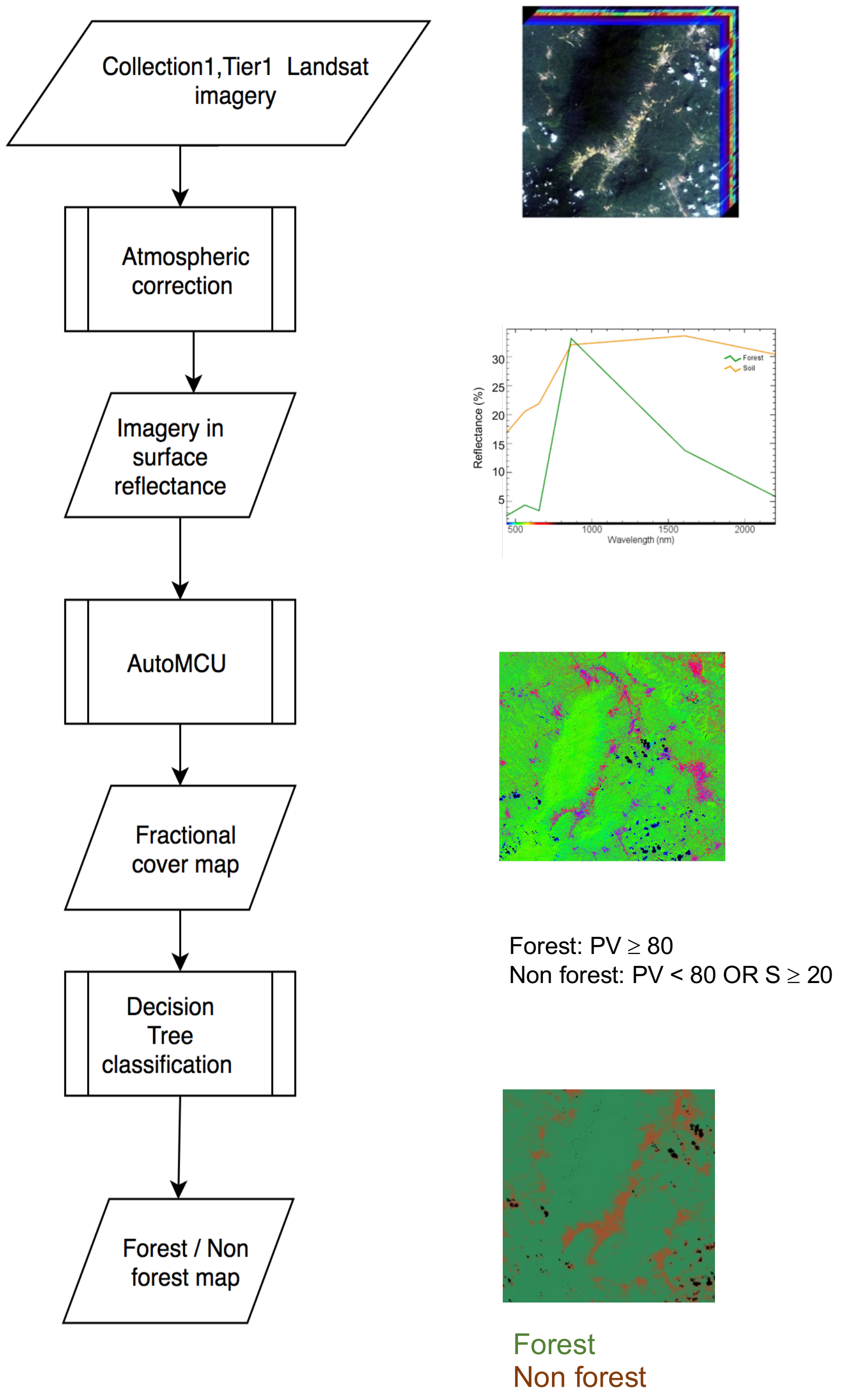
Collection1,Tier1 Landsat imagery
Atmospheric correction
Imagery in surface reflectance
AutoMCU
Fractional cover map
Decision Tree classification
Forest / Non forest map
Reflectance (%)
Wavelength (nm)
Forest
Soil
Forest: PV ≥ 80
Non forest: PV < 80 OR S ≥ 20
Forest
Non forest

B

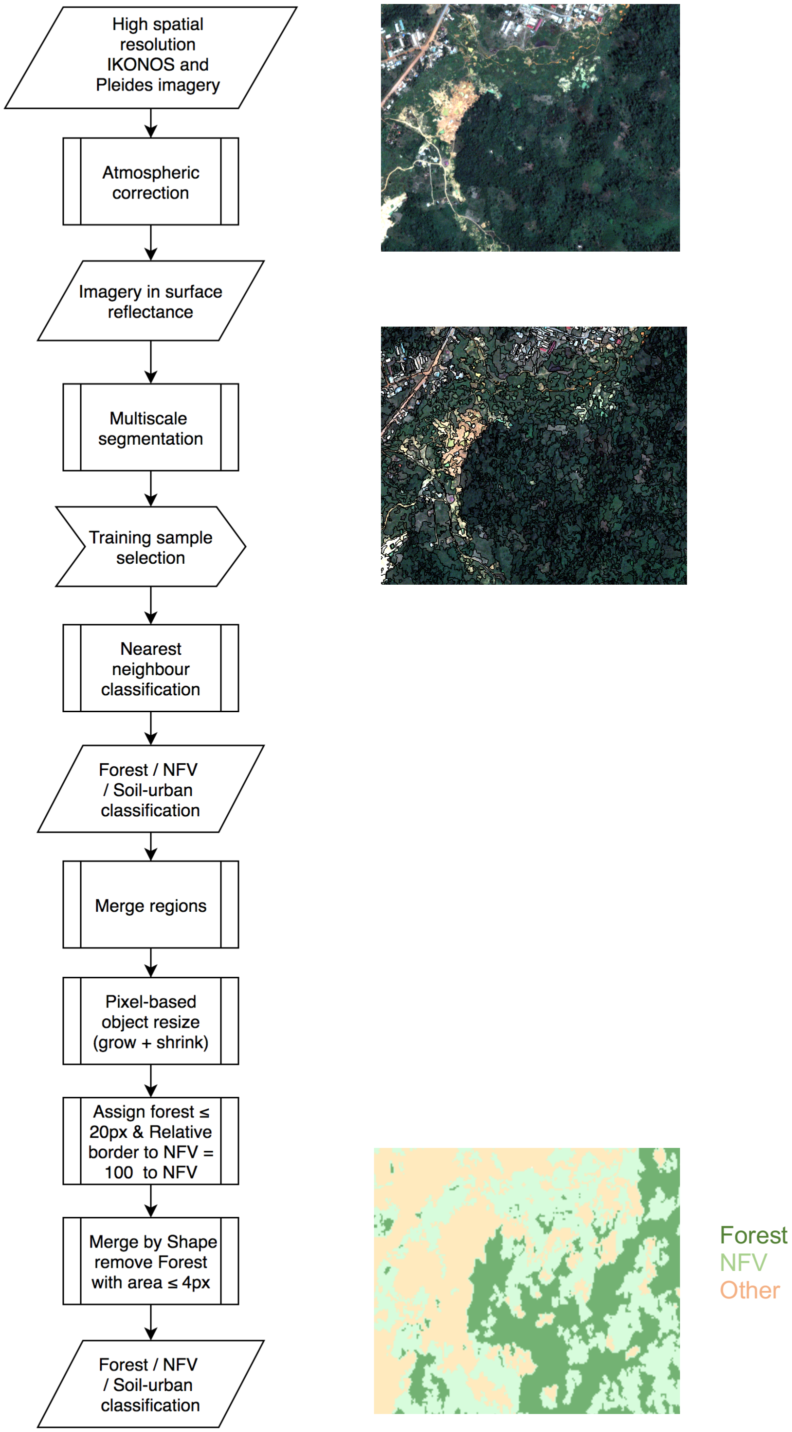


FIGURE S5 A) Flowchart of the analytical steps to generate the forest / non forest classification for each Landsat image. Examples to the right of the flowchart are from top to bottom: Landsat image cube, spectral signatures from forest and soil pixels (in surface reflectance), fractional cover map with non-photosynthetic vegetation (NPV)/Photosynthetic vegetation (PV) / Bare substrate (S) as a NPV:PV:S composite and final classification map with forest and non-forest classes. All steps were carried out in CLASlite v 3.2. B) Flowchart of the main analytical steps to generate the classification for the high-resolution satellite imagery. Examples to the right of the flowchart are from top to bottom: an atmospherically corrected Pleides image, the same image having undergone segmentation – the black polygons outline each segment, final classification for forest, non forest vegetation (NFV) and other (including soil, water, built up areas). Atmospheric correction was carried out in ENVI 5.5. All other steps were carried out in eCognition Developer 9.4.

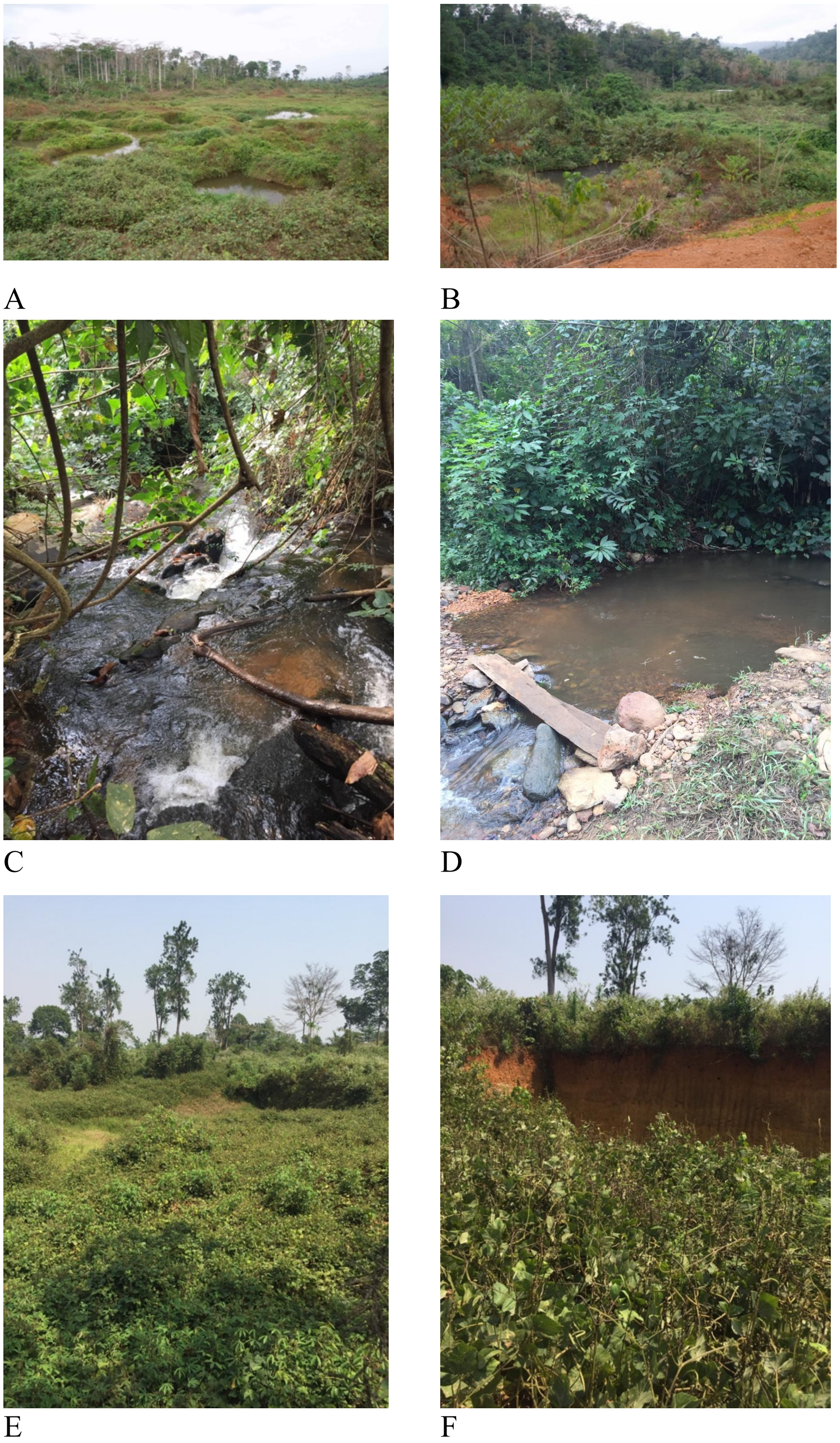


Figure S6. a,b) Photographs from Feb 2018 of the location of former streams located approximately halfway between Kibi and Asiakwa; c,d)Photographs collected Jan 2019 from creeks near the type locality (yellow points on S7); e) herbaceous vegetation growth near the type locality (Jan 2019); f) abandoned ASGM pit at the coordinates of the type locality.

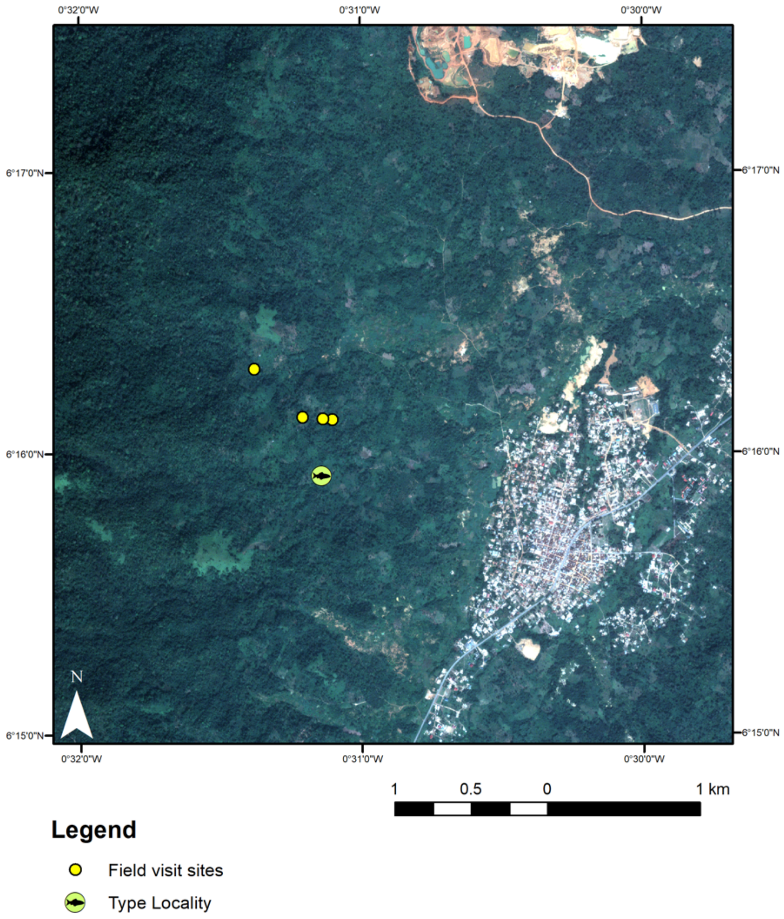


Figure S7. Pleides 1B image acquired Dec 9, 2018 for the type locality of *L. robertsi*, west of Asiakwa. Yellow points represent small higher altitude creeks visited in Jan 2019.

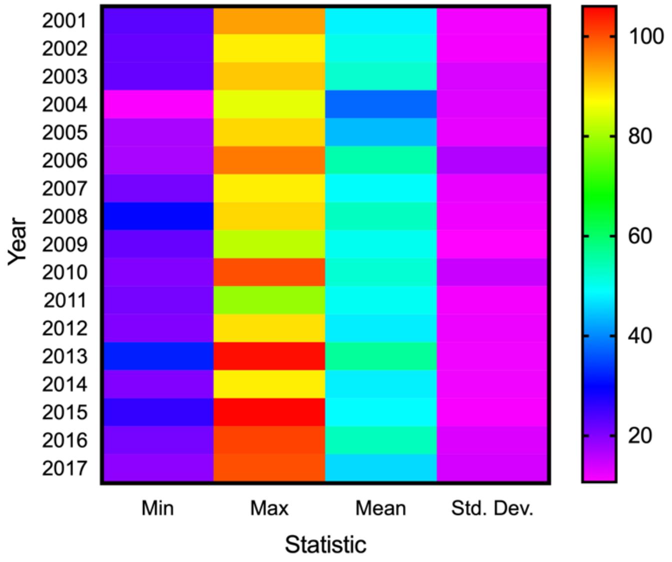


Figure S8. Yearly cloud cover statistics for the study area extracted from the MODIS/Terra Surface Reflectance Daily L2G Global 1 km SIN Grid V006 MOD09GA. Values represent number of cloud free days per year (1 km pixels).
